# Deep motif deconvolution of HLA-II peptidomes for robust class II epitope predictions

**DOI:** 10.1101/539338

**Authors:** Julien Racle, Justine Michaux, Georg Alexander Rockinger, Marion Arnaud, Sara Bobisse, Chloe Chong, Philippe Guillaume, George Coukos, Alexandre Harari, Camilla Jandus, Michal Bassani-Sternberg, David Gfeller

## Abstract

CD4 T cells are key for priming and regulating immune recognition of infected and cancer cells, but predictions of class II epitopes have limited accuracy. We combined unbiased Mass Spectrometry-based HLA-II peptidomics with a novel motif deconvolution algorithm to profile and analyze a total of 99’265 unique HLA-II ligands. Our work demonstrates substantial improvement in the definition of HLA-II binding motifs and enhanced accuracy in class II epitope predictions.

## Main

Antigen presenting cells (APCs) display on their surface peptides bound to class II Human Leukocyte Antigen molecules (HLA-II). In the case of infections or cancer, presentation by APCs and recognition by CD4+ T cells of non-self-peptides or tumor-associated antigens are key to initiate and sustain an immune response^1–4^. The binding of peptides to their cognate HLA-II alleles is thus of fundamental importance. However, analysis of binding specificity of HLA-II alleles is challenging due to their high polymorphism and the open binding groove that allows peptides of diverse sizes to bind. This makes alignment of HLA-II ligands difficult, especially in HLA peptidomics data where peptides found in a sample come from multiple alleles. Accordingly, class II epitope predictions display low accuracy^5,6^. Recent developments in HLA peptidomics^7–9^ are promising to improve epitope predictions, as shown in multiple studies of HLA-I molecules^10–14^. However, similar improvements in class II epitope predictions have not been observed and early studies have been restricted to very few (1 to 3) HLA-II alleles^8,15^.

Here, we profiled the HLA-II peptidome of 13 different cell lines or tissue samples and identified 40’864 unique HLA-II ligands. We completed these data with our recent publication^7^ to reach a total of 77’189 unique peptides from 23 different cell lines or tissue samples (see Methods, Fig. 1a, Suppl. Table 1 and Suppl. Data 1), making it the largest dataset of HLA-II ligands available to date. To analyze these data, we developed MoDec, a novel Motif Deconvolution algorithm (see Methods). Unlike previous approaches to analyze HLA-II ligands that attempted to align the peptides^16,17^, MoDec is a fully probabilistic framework that allows motifs to be found everywhere on the peptide sequences and learns both the motifs as well as their respective weights and preferred binding core position offsets (Fig. 1b). MoDec shows conceptual similarity with convolutional neural networks (each motif can be thought of as a filter) but provides direct interpretation and visualization of the results as sequence logos (Fig. 1b). Applying MoDec to our data, we found an abundance of motifs (Suppl. Fig. 1). HLA-II binding motifs identified across samples with shared HLA-II alleles displayed very high similarity (Fig. 1c and Suppl. Fig. 2). This demonstrates high reproducibility of our motif deconvolution approach and enabled us to unambiguously annotate the different motifs to their respective alleles. Comparison with known motifs from databases such as IEDB^18^, or predictions from NetMHCIIpan^19^ showed some similarity but also some important differences (Suppl. Fig. 2). In particular, the motifs deconvolved from HLA-II peptidomics data tend to show clearer anchor residues and some motifs displayed a novel type of binding specificity (Suppl. Fig. 2).

**Figure 1.**
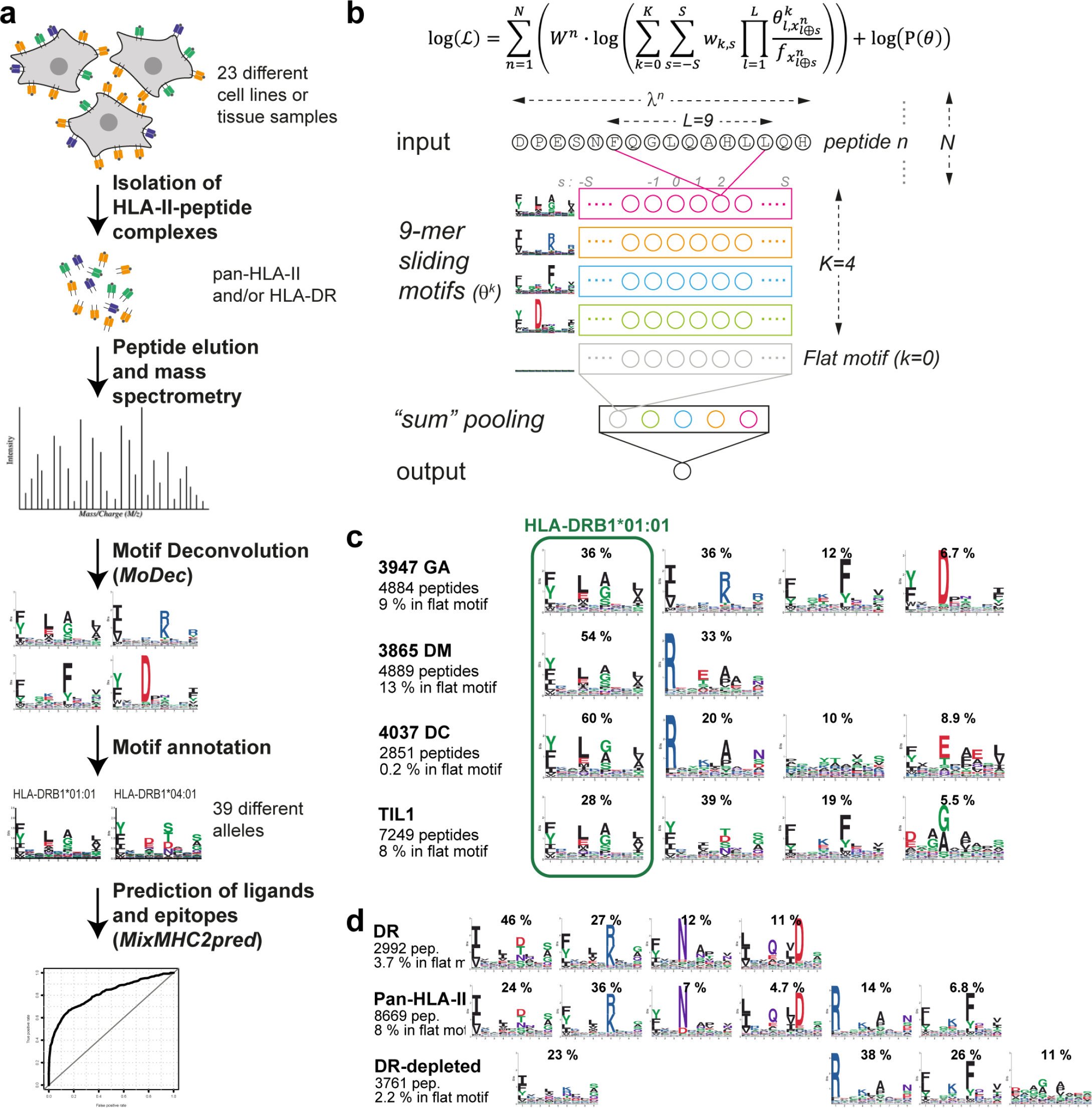
Motif deconvolution in HLA-II peptidomics data. **(a)** Description of our pipeline for MS-based HLA-II ligand isolation, motif deconvolution with MoDec and training of an HLA-II ligand predictor. **(b)** Log-likelihood optimized in MoDec (see Methods). Below is a graphical interpretation of the model, including multiple sliding motifs of 9 amino acids (*θ*^*k*^) and a sum pooling step over all possible positions (*s*) of the 9-mer motifs. **(c)** Motifs identified in four samples sharing exactly one allele (HLA-DRB1*01:01) and showing exactly one highly conserved motif. **(d)** Comparison of the motifs found in HLA-II peptidomics, HLA-DR peptidomics and HLA-DR-depleted peptidomics in the same tissue sample (*3830 NJF*).

**Figure 2.**
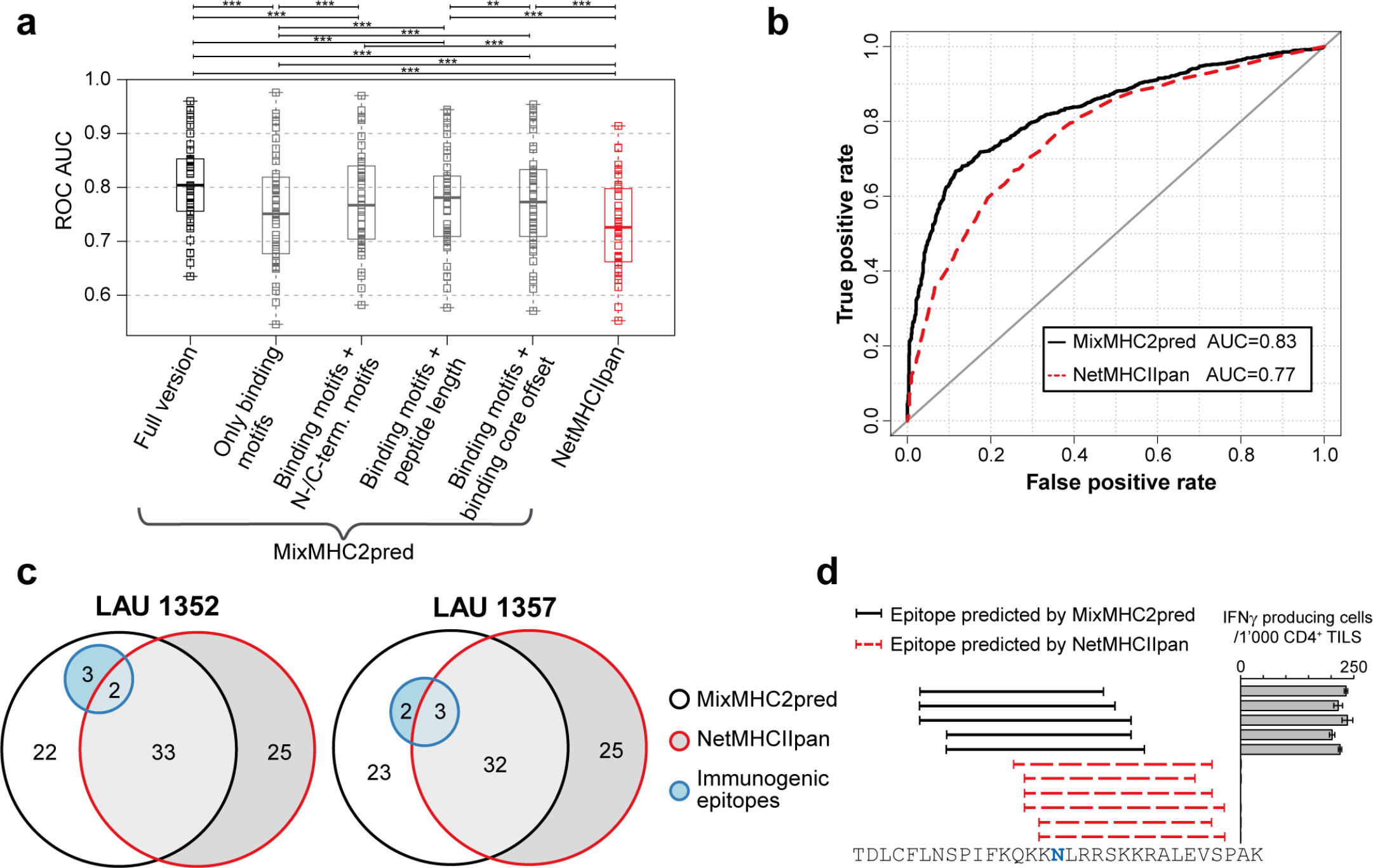
MixMHC2pred improves the prediction accuracy for HLA-II ligands and class II epitopes. **(a)** Comparison of prediction accuracy of MixMHC2pred (and multiple variants) and NetMHCIIpan for HLA-II ligands. Paired Wilcoxon signed rank test is performed (**: p-value < 0.01, ***: p-value < 0.001). **(b)** ROC curve for the predictions of all class II epitopes from CD4+ T cell tetramer assays in IEDB^18^. **(c)** Euler diagram showing the number of potential epitopes from viral, bacterial and melanoma-associated antigens that were tested based on MixMHC2pred and NetMHCIIpan predictions and how many of these were truly immunogenic (blue circles) in two different melanoma patients. **(d)** Predictions with MixMHC2pred and NetMHCIIpan of candidate epitopes in a minigene encoding an ovarian cancer class II immunogenic mutation (SGOL1_*D246N*_). The bar plot shows the resulting CD4+ T cell response towards these epitopes.

The clearest and most frequent motifs that we identified correspond to HLA-DR alleles. To further validate them, we sequentially purified first the HLA-DR molecules with anti-HLA-DR antibody and then the remaining HLA-II molecules with a pan-HLA-II antibody (42’903 and 27’692 unique peptides respectively, leading to a total of 99’265 unique peptides across all our samples, see Methods and Suppl. Data 2). We observed that the motifs deconvolved from HLA-DR peptidomes are identical to those assigned to HLA-DR alleles from the pan-HLA-II peptidomes (Fig. 1d and Suppl. Fig. 3). In addition, in the HLA-DR-depleted samples, all motifs previously predicted to correspond to HLA-DP or - DQ alleles could be found, indicating that our motif deconvolution approach in pan-HLA-II peptidomes is not restricted to HLA-DR motifs and that MoDec performs well in pan– HLA-II data (Fig. 1d and Suppl. Fig. 3). Compared to other motif deconvolution methods^17,20^, MoDec shows improved resolution and is 100-10’000 times faster (Suppl. Fig. 4), making it particularly well suited for deconvolution in large HLA-II peptidomics samples.

We next investigated properties of naturally presented HLA-II ligands beyond the binding specificity of HLA-II alleles. We first grouped all our HLA-II ligands and used MoDec to identify motifs occurring three amino acids upstream and downstream of the N- or C-terminus of the peptides (see Methods). Multiple motifs appeared (Suppl. Fig. 5a-b), some of which had been suggested to represent different peptide processing and cleavage pathways^21^. The specificity was in general stronger for amino acids inside the peptide than those in flanking regions (Suppl. Fig. 5a-b). In particular, clear enrichment was observed for proline two residues downstream of the N-terminus and upstream of the C-terminus, confirming previous observations made for a limited number of alleles^15,21^. All alleles showed such a proline enrichment (Suppl. Fig. 5c-d). Binding assay experiments for three HLA-DRB1 alleles demonstrate that the binding affinity is similar between peptides containing proline or alanine at the 2^nd^ position (Suppl. Fig. 5e), validating the hypothesis that proline enrichment is due to the peptide processing and loading independently of the binding specificity of the HLA-II molecules. We then observed that the peptide length distribution was conserved between HLA-II alleles (Suppl. Fig. 6a), unlike for HLA-I alleles^22^. The distribution of binding core offsets was also independent of the alleles (Suppl. Fig. 6b), with a slight shift of the core towards the C-terminus (i.e. slightly more amino acids extend outside of the binding core on the N-terminus than the C-terminus).

We then took advantage of our deconvolved HLA-II peptidomics datasets to train a predictor of HLA-II ligands (MixMHC2pred, see Methods). This predictor combines allele-specific peptide binding motifs and allele-independent peptide N-/C-terminal motifs, peptide length and binding core offset preferences (see Methods). We first performed predictions on multiple HLA-II peptidomics datasets from independent studies (Suppl. Table 2) and observed improved accuracy with our predictor compared to NetMHCIIpan^19^, the state-of-the-art predictor for HLA-II ligands (Fig. 2a, Suppl. Fig. 7). These results also show the superiority of the full predictor combining the binding motifs, N-/C-terminal motifs, peptide length and binding core offset, compared to a predictor based only on the binding motifs or a combination of the binding motifs with any of the other allele-independent characteristics (Fig. 2a, Suppl. Fig. 7). To investigate whether MixMHC2pred was appropriate not only for HLA-II ligands but also for class II epitope predictions, we compiled all CD4+ T cell tetramer assays found in IEDB^18^. Receiver operating characteristic (ROC) curve analysis showed that MixMHC2pred was more accurate than NetMHCIIpan (Fig. 2b). We further surveyed a set of several known melanoma-associated antigens and viral and bacterial proteins (Suppl. Table 3a and Methods). Top hits from MixMHC2pred and from NetMHCIIpan for tumor-associated antigens (top 30 peptides) and viral/bacterial antigens (top 30 peptides) were tested for class II immunogenicity in two melanoma patients (Suppl. Table 3b and Methods). Our results show an enrichment for immunogenic epitopes among the predictions of MixMHC2pred versus NetMHCIIpan (Fig. 2c, Matthews correlation coefficients of 0.16 and 0.16 for MixMHC2pred versus −0.17 and −0.058 for NetMHCIIpan). The same peptides were also tested with CD4+ T cells from a healthy donor resulting in many correctly predicted epitopes and similar yield between the two predictors (Suppl. Fig. 8a). We next took advantage of a class II immunogenic mutation (*D246N* in SGOL1) that was recently identified in an ovarian cancer patient by screening tumor infiltrating lymphocytes (TILs) with minigenes (see Methods and Suppl. Fig. 9). To determine the actual epitope, we applied both MixMHC2pred and NetMHCIIpan on the 31-mer coded by the minigene initially used to test immunogenicity. The results indicate that here again our approach could accurately predict the actual epitope from the 31-mer (Fig. 2d, Suppl. Fig. 10 and Methods).

By combining in-depth HLA-II peptidomics data of 49 different samples together with a novel motif deconvolution approach (MoDec), we show that we can capitalize on accurate and unbiased MS profiling of HLA-II ligands for class II epitope predictions. The very high similarity between HLA-DR motifs identified in pan-HLA-II and HLA-DR peptidomes shows that all HLA-DR motifs can be accurately resolved in the pan-HLA-II samples with MoDec. This indicates that mono-allelic samples are not needed to determine HLA-DR motifs, thereby greatly reducing the amount of experimental work of future studies. Whether increased detection efficacy could be obtained by using anti-HLA-DP and anti-HLA-DQ antibodies remains to be seen and our motif deconvolution tool may prove highly valuable to analyze such data once they become available. Although our approach may still not capture the full complexity of class II antigen presentation (e.g., differences between endocytic and phagocytic pathways^1^), the use of MS data enabled us to incorporate N- and C-terminal motifs, as well as peptide length and binding core offset preferences, in addition to binding specificity of HLA-II alleles. The large allele coverage (especially for HLA-DR alleles) makes our predictor suitable for a wide range of applications in infectious diseases, autoimmunity and cancer immunotherapy.

## Methods

Methods and any associated references are available in the online version of the paper.

## Acknowledgments

We thank the Center of Experimental Therapeutics team for providing us with the patient-derived tissue samples and T cells. We are thankful to Pedro Romero for sharing the B-cell lines with us. We thank Roy T. Daniel from the Service of Neurosurgery at the University Hospital of Lausanne, and Monika Hegi for providing us with the collection of meningioma tissues. We thank Marthe Solleder for help with the visualization of motifs with ggseqlogo, Fabio Marino for technical support with sample preparation, HuiSong Pak for MS measurements and Raphaël Genolet for HLA typing.

This work was supported by the Swiss Cancer League (KFS-4104-02-2017, DG and JR), by the Ludwig Institute for Cancer Research, by the ISREC Foundation thanks to a donation from the Biltema Foundation (JM, CC and MBS) and by the MEDIC foundation (GAR and CJ).

## Author contributions

JR developed the computational methods; JR and DG analyzed the data; JM, CC and MBS generated the MS peptidomics data; GAR, MA, SB, PG, AH and CJ performed the binding and T cell assays; GC, AH, CJ and MBS provided reagents; JR, MBS and DG designed the study; JR, MBS and DG wrote the paper.

## Online Methods

### Cells and patient material

EBV-transformed human B-cell lines JY (ATCC^®^ 77442™, Manassas, Virginia), CD165, PD42, CM467, RA957, BP455, GD149 (a gift from Pedro Romero, Ludwig Cancer Research Lausanne), were maintained in RPMI 1640 + GlutaMAX medium (Life Technologies, Carlsbad, California) supplemented with 10% heat-inactivated FBS (Dominique Dutscher, Brumath, France) and 1% Penicillin/Streptomycin Solution (BioConcept, San Diego, California). Cells were grown to the required cell amount, collected by centrifugation at 1,200 rpm for 5 min, washed twice with ice cold PBS and stored as dry cell pellets at −20°C until use.

T-cells were expanded from two melanoma tumors as previously described^7,11^ following established protocols^23,24^. Briefly, fresh tumor samples were cut in small fragments and placed in 24-well plates containing RPMI CTS grade (Life Technologies), 10% Human serum (Valley Biomedical, Winchester, Virginia), 0.025 M HEPES (Life Technologies), 55 μmol/L 2-Mercaptoethanol (Life Technologies) and supplemented with a high concentration of IL-2 (Proleukin, 6,000 IU/mL, Novartis, Basel, Switzerland) for three to five weeks. 25 ×10^6^ TIL were stimulated with irradiated feeder cells, anti-CD3 (OKT3, 30 ng/mL, Miltenyi biotec) and high dose IL-2 (3,000 IU/mL) for 14 days. The cells were washed using a cell harvester (LoVo, Fresenius Kabi, Lake County, Illinois). Finally, the cells were washed with PBS on ice, aliquoted to a cell count of 1 × 10^8^ and stored as dry pellets at −80°C until use.

Snap frozen meningioma tissues from patients (3808-HMC, 3830-NJF, 3849-BR, 3865-DM, 3869-GA, 3911-ME, 3912-BAM, 3947-GA, 3971-ORA, 3993, 4001, 4021, 4037-DC, 4052-BA) were obtained from the Centre hospitalier universitaire vaudois (CHUV, Lausanne, Switzerland). Informed consent of the participants was obtained following requirements of the institutional review board (Ethics Commission, CHUV). Protocol F-25/99 has been approved by the local Ethics committee and the biobank of the Lab of Brain Tumor Biology and Genetics. Protocol 2017-00305 for Antigens and T cells discovery in tumors has been approved by the local Ethics committee.

### HLA typing

Genomic DNA was extracted using DNeasy kit from Qiagen and 500 ng of gDNA were used to amplify HLA genes by PCR. High resolution 4-digit HLA typing was performed with the TruSight HLA v2 Sequencing Panel from Illumina on a MiniSeq instrument (Illumina). Sequencing data were analyzed with the Assign TruSight HLA v2.1 software (Illumina) and are provided as Suppl. Table 1.

### Generation of antibody-crosslinked beads

Anti-pan-HLA-II and anti-HLA-DR monoclonal antibodies were purified from the supernatant of HB145 (ATCC^®^ HB-145™) and HB298 cells (ATCC^®^ HB-298™), respectively, grown in CELLLine CL-1000 flasks (Sigma-Aldrich, St. Louis, Missouri) using Protein A-Sepharose 4B beads (Pro-A beads; Invitrogen, Carlsbad, California). Antibodies were cross-linked to Pro-A beads at a concentration of 5 mg of antibodies per 1 mL volume of beads. For this purpose, the antibodies were incubated with the Pro-A beads for 1 hour at room temperature. Chemical cross-linking was performed by addition of Dimethyl pimelimidate dihydrochloride (Sigma-Aldrich) in 0.2 M Sodium Borate buffer pH 9 (Sigma-Aldrich) at a final concentration of 20 mM for 30 minutes. The reaction was quenched by incubation with 0.2 M ethanolamine pH 8 (Sigma-Aldrich) for 2 hours. Cross-linked antibodies were kept at 4°C until use.

### Purification of HLA-II and HLA-DR peptides

Cells were lysed in PBS containing 0.25% sodium deoxycholate (Sigma-Aldrich), 0.2 mM iodoacetamide (Sigma-Aldrich), 1 mM EDTA, 1:200 Protease Inhibitors Cocktail (Sigma-Aldrich), 1 mM Phenylmethylsulfonylfluoride (Roche, Basel, Switzerland), 1% octyl-beta-D glucopyranoside (Sigma-Aldrich) at 4°C for 1 hour. The lysis buffer was added to the cells at a concentration of 1 × 10^8^ cells/mL. Cell lysates were cleared by centrifugation with a table-top centrifuge (Eppendorf Centrifuge, Hamburg, Germany) at 4°C at 14,200 rpm for 50 min. Meningioma tissues were placed in tubes containing the same lysis buffer and homogenized on ice in 3-5 short intervals of 5 seconds each using an Ultra Turrax homogenizer (IKA, Staufen, Germany) at maximum speed. For one gram of tissue, 10 to 12 mL of lysis buffer was required. Cell lysis was performed at 4°C for 1 hour. Tissue lysates were cleared by centrifugation at 20,000 rpm in a high-speed centrifuge (Beckman Coulter, JSS15314, Nyon, Switzerland) at 4°C for 50 minutes. The cells and tissue lysates were loaded on stacked 96-well single-use micro-plates (3μm glass fiber, 10μm polypropylene membranes; ref number 360063, Seahorse Bioscience, North Billerica, Massachusetts). Purification of pan-HLA-II peptides were performed following depletion of HLA-I as previously described^7^. For the sequential purification of HLA-DR and HLA-II from tissues, three plates were used. The first plate contained protein-A sepharose 4B (Pro-A) beads (Invitrogen, Carlsbad, California) for depletion of antibodies (pre-clear plate), the second plate contained same beads cross-linked to the anti HLA-DR monoclonal antibodies and the third plate contained the beads cross-linked to anti HLA-II monoclonal antibodies. For the sequential purification of HLA-DR and HLA-II from cells, only the last two plates were used. The Waters Positive Pressure-96 Processor (Waters, Milford, Massachusetts) was employed. The second and third plates were washed separately with 4 times 2 mL of 150 mM sodium chloride (NaCl) (Carlo-Erba, Val de Reuil, France) in 20 mM Tris-HCl pH 8, 4 times 2 mL of 400 mM NaCl in 20 mM Tris-HCl pH 8 and again with 4 times 2 mL of 150 mM NaCl in 20 mM Tris-HCl pH 8. Finally, the plates were washed twice with 2 mL of 20 mM Tris-HCl pH 8. Each affinity plate was stacked on top of a Sep-Pak tC18 100 mg Sorbent 96-well plate (ref number: 186002321, Waters) already equilibrated with 1 mL of 80% acetonitrile (ACN) in 0.1 % TFA and with 2 mL of 0.1% TFA. The HLA and peptides were eluted with 500 μL 1% TFA into the Sep-Pak plate and then we washed this plate with 2 mL of 0.1 % TFA. Thereafter, HLA-II and HLA-DR peptides were eluted with 500 μL of 32% ACN in 0.1% TFA into a collection plate. Recovered HLA-II and HLA-DR peptides were dried using vacuum centrifugation (Concentrator plus Eppendorf) and stored at −20°C.

### LC-MS/MS analyses of HLA-II peptides

Prior to MS analysis HLA-II and HLA-DR peptide samples were re-suspended in 10 μL of 2% ACN in 0.1 % FA and aliquots of 3 uL for each MS run were placed in the Ultra HPLC autosampler. HLA peptides were separated by a nanoflow HPLC (Proxeon Biosystems, Thermo Fisher Scientific, Odense) coupled on-line to a Q Exactive HF or HFX mass spectrometers (Thermo Fisher Scientific, Bremen) with a nanoelectrospray ion source (Proxeon Biosystems). We packed a 20 cm long, 75 μm inner diameter column with ReproSil-Pur C18-AQ 1.9 μm resin (Dr. Maisch GmbH, Ammerbuch-Entringen, Germany) in buffer A (0.5% acetic acid). Peptides were eluted with a linear gradient of 2–30% buffer B (80% ACN and 0.5% acetic acid) at a flow rate of 250 nl/min over 90 min. Data was acquired using a data-dependent ‘top 10’ method. Full scan MS spectra were acquired at a resolution of 70,000 at 200 m/z with an Auto gain control (AGC) target value of 3e6 ions. Ten most abundant ions were sequentially isolated, activated by Higher-energy Collisional Dissociation and accumulated to an AGC target value of 1e5 with a maximum injection time of 120 ms. In case of assigned precursor ion charge states of one, and from six and above, no fragmentation was performed. MS/MS resolution was set to 17,500 at 200 m/z. Selected ions from Ions were dynamically excluded for additional fragmentation for 20 seconds. The peptide match option was disabled. The raw files and MaxQuant output tables will be available with the peer reviewed version of this manuscript.

### Peptide identification

We employed the MaxQuant platform^26^ version 1.5.5.1 to search the peak lists against a fasta file containing the human proteome (Homo_sapiens_UP000005640_9606, the reviewed part of UniProt, with no isoforms, including 21,026 entries downloaded in March 2017) and a list of 247 frequently observed contaminants. Peptides with a length between 8 and 25 AA were allowed. The second peptide identification option in Andromeda was enabled. The enzyme specificity was set as unspecific. A false discovery rate (FDR) of 1% was required for peptides and no protein FDR was set. The initial allowed mass deviation of the precursor ion was set to 6 ppm and the maximum fragment mass deviation was set to 20 ppm. Methionine oxidation and N-terminal acetylation were set as variable modifications. The Peptide output files summarizing MaxQuant result files are provided as Supplementary Data 1 and 2.

### Motif deconvolution algorithm for HLA-II peptidomics

In HLA-II peptidomics data, the HLA-II ligands are coming from different alleles, are of different lengths, and their binding core positions are a priori unknown. To account for this, and building upon the successful application of the mixture model to HLA-I peptidomics^10^, we developed a probabilistic framework able to learn multiple motifs anywhere on the peptides, as well as the weights and binding core offsets of these motifs. The log-likelihood is given by the following equation (see also Fig. 1b):

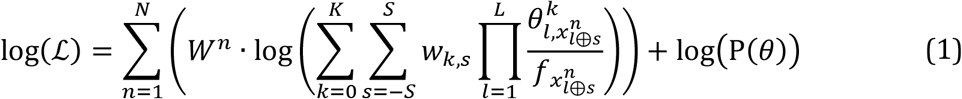

where *N* is the number of peptides; *W*^*n*^ is the similarity weight of *n*^th^ peptide (see below); *K* is the number of motifs; *S* is the maximal binding core offset 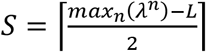, where *λ*^*n*^ is the length of *n*^th^ peptide); *w*_*k,s*_ is the weight of motif *k* with binding core offset *s* (∑_*k*_ ∑_*s*_ *w*_*k*,*s*_ = 1); *L* is the binding motif length (equal to 9 here since HLA-II ligands are known to bind with a 9-mer core); *θ*^*k*^_*l,i*_ represent the binding motifs (with 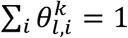; *k=0* is a special case of a flat motif: 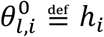, where *h*_*i*_ are the amino acid frequencies in the human proteome (this motif is used to model potential contaminant peptides)); *x*^*n*^_*j*_ indicates which amino acid is found in peptide *n* at the position *j* (when *x*^*n*^_*j*_ is not defined (i.e. *j<1* or *j>λ^n^*), then 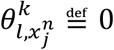; *f*_*i*_ is the expected background frequency in HLA-II peptidomics data for amino acid *i*; and P(*θ*) is a Dirichlet prior term (with the hyper-parameter equal to 0.1)^27^. The “*l* ⊕ *s*” in 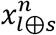 is a “special sum” that makes that the binding core offsets are symmetric around 0 for each peptide (see Supplementary Note). A peptide similarity weight is given by 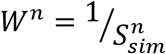, where *S*^*n*^_*sim*_ represents the average number of times each 9-mer from the *n*^*th*^ peptide is observed in the full dataset. This is useful since multiple overlapping HLA-II ligands from the same source region are typically found by MS.

Unlike the previous approaches for HLA class I^10,11^, our model does not need previous peptide alignment, learns the binding core offsets of the motifs and includes peptide similarity weights. MoDec estimates the parameters *θ* and *w* based on expectation-maximization (see Suppl. Note for details). Although our framework is fully probabilistic, peptide responsibilities are derived during the expectation-maximization, and these can be used to predict to which motif each peptide is most likely associated and with which binding core offset. Multiple runs (250 in this study) are performed by MoDec to optimize the log-likelihood of equation (1), starting from different initial conditions, considering all peptides of length 12 or more. As HLA-DR ligands have preference for hydrophobic amino acids at position 1, we implemented the possibility to include such a bias in a subset of the initial conditions used in the optimization by MoDec.

Binding motifs determined by MoDec are visualized with ggseqlogo^28^.

The optimal number of motifs (*K*) was first determined using the Akaike Information Criterion (AIC):

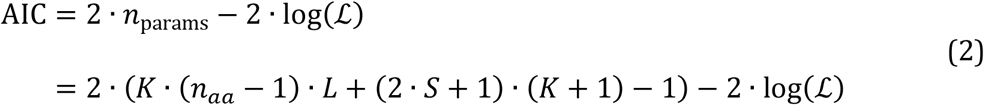

where *n*_*params*_ is the number of free parameters, *n*_*aa*_ is the number of different amino acids (20) and the other parameters have been defined earlier. This Akaike Information criterion is commonly used in information theory to determine the information gained from using a model with more parameters over a simpler model (the smallest AIC value the better). However, as with HLA-I peptidomics data^10^ (and more generally with many clustering approaches), the optimal number of motifs is difficult to determine in a fully unsupervised way. We therefore explored additional motifs, tried further splitting specific motifs, and manually curated each dataset for consistency across samples. Comparison between the numbers of motifs manually curated or determined by the AIC shows that in most cases the correct number of motifs would have been found with AIC and in 80% of the cases the error when using this criterion would be at most of 1 motif (Supplementary Fig. 11), suggesting that AIC is a good starting point for the selection of the optimal number of motifs.

### Assignment of motifs to alleles

To annotate the different motifs to their respective alleles, we used an iterative approach: we considered all samples that share a given allele and determined if a motif was shared between all these samples. To decide which motifs are shared, we used the Kullback-Leibler divergence between the motifs 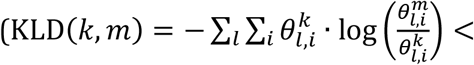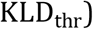 for a given threshold (*KLD*_*thr*_ comprised between 1 and 1.75 depending on the iteration). Each iteration consists in checking for each allele, one after the other, if we can assign a motif to this allele. In the first 5 iterations, at least 75% of the samples containing the given allele had to share a motif in order to assign this motif to the given allele (in later iterations, the threshold of samples is decreased to 60%), with the additional requirement that no other allele was shared by these samples. By repeating these iterations multiple times, various motifs could be annotated to their respective allele. All annotations were further manually curated, allowing for example the annotation of HLA-DRB1*01:02 (that is highly similar to HLA-DRB1*01:01 but with only the *PD42* sample expressing this allele – see Suppl. Fig. 2).

For the binding motifs from IEDB^18^ database, we downloaded the full MHC ligand data (http://www.iedb.org/database_export_v3.php, version from 28.01.2018) and filtered this data to remove peptides obtained from mass spectrometry (since for many of them allele restriction information is based on predictions) and to keep only the peptides described as “Positive-High” binders. MoDec was run considering a single motif on the resulting list of peptides per allele. The corresponding motifs are shown in Suppl. Fig. 2.

Binding motifs from NetMHCIIpan^19^ were determined in the following way: 16’000 random human peptides (2’000 of each length between 12 and 19 amino acids) were inputted into NetMHCIIpan. For each allele, the peptides with a “%Rank” better than 5 were kept and ggseqlogo^28^ was used to draw the motifs from the corresponding “Core” sequences returned by NetMHCIIpan.

### Comparison to other motif deconvolution methods

We compared the motifs found by MoDec with those predicted by Gibbscluster (version 2.0^17^) and MEME version 4.12^20^ (Suppl. Fig. 4). For a comparison of the timing, the tools were launched on a single 3.3 GHz CPU with 2 GB of RAM, searching for 1 to 8 motifs in various samples.

Gibbscluster was run with the recommended parameters for HLA-II (5 seeds for the initial conditions, an initial MC temperature of 1.5, using a trash cluster with a threshold of 2 for this cluster, the rest being left unchanged). Gibbscluster suggests using a Kullback-Leibler criterion to select the optimal number of clusters from their deconvolution. In some cases, manual curation allowed to find additional clusters that had similarity with known HLA-II motifs, and we show the results of Gibbscluster including these additional clusters in Suppl. Figure 4.

MEME was run setting a motif width of 9, a maximum dataset size of 10’000’000 (needed due to the size of the samples) and the rest was left at default.

### Investigation of properties of HLA-II ligands other than binding specificity

Analysis of N- and C-terminal flanking motifs was done with MoDec by taking the three amino acids upstream and downstream of the N-/C-terminus of the peptides that could be assigned to alleles (i.e., the peptides were extended based on their protein of origin to include the three amino acids upstream of N-terminus and downstream of C-terminus). The peptide length distributions, binding core offset distributions and frequencies of proline two residues downstream of the N-terminus and upstream of the C-terminus, were computed for all peptides associated to each allele.

### Binding affinity assays

Peptide binding affinity (Suppl. Fig. 5e) was assessed by peptide competition assay. For each peptide, eight wells of a v-bottom 96-well plate (Greiner Bio-One) were filled with 100 μl of each recombinant “empty” DR1, DR4 or DR7 protein (1 μg) in a citrate saline buffer (100 mM citrate pH 6.0), with 0.2% β-octyl-glucopyranoside (Calbiochem), 1xcomplete protease inhibitors (Roche) and 2 μM FLAG-HA307-319 peptide. Competitor peptides (10 mM DMSO solution) were added to each well to a final concentration of either 100, 33, 11, 3.7, 1.2, 0.4, 0.1 and 0 μM for DR1 and DR4 or 100, 33, 11, 3.7, 1.2, 0.4, 0.1 and 0 nM for DR7. After incubation at 37°C overnight, 100 μl was transferred to a plate coated with avidin (2 μg/ml) and previously blocked. After 1 h of incubation at RT and three washes with 1×PBS pH 7.4, 0.05% Tween 20, anti-FLAG-alkaline phosphatase conjugate (Sigma) was added as 1:5,000. After 1 h, the plate was washed as previously described and developed with pNPP SigmaFAST substrate and absorbance read with the 405 nm filter.

### MixMHC2pred – a predictor of HLA-II ligands

We trained a predictor of HLA-II ligands (*MixMHC2pred*) using all our HLA-II peptidomics data (including the pan-HLA-II, HLA-DR and HLA-DR depleted peptidomics data). For a given allele, *a*, and peptide, *n*, the binding score is given by:

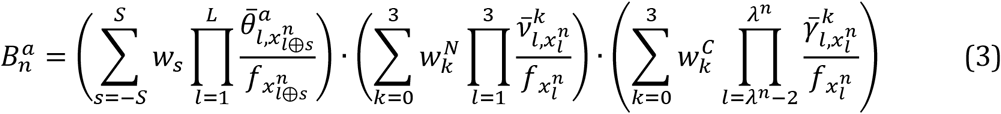

where *w*_*s*_ represents the global binding core offset preference (computed combining all peptides associated to an allele); 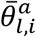 is the position probability matrix for allele *a* (computed from all peptides associated to this allele with their respective binding core offset based on the highest responsibility value, and adding pseudocounts based on the BLOSUM62 substitution matrix with a parameter *β*=200^29^); 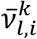 and 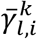 are similar matrices representing the N- and C-terminal motifs (Suppl. Fig. 5a-b; including here only the amino acids within the peptides); *w*^*N*^_*k*_ and *w*^*C*^_*k*_ represent the relative contributions of the N-/C-terminal motifs (i.e. the fraction of peptides assigned to each of these motifs). See Equation (1) for the definition of other terms.

This binding score is then transformed to a percentile rank per peptide length by comparing it to the score of 10’000 random human peptides of the same length, and then further transformed to a global percentile rank by making that the top 1% random human peptides follow the same peptide length distribution than the global peptide length distribution observed in our HLA-II peptidomics data. Finally, when the score among multiple alleles is requested, the score from each peptide is taken as its best percentile rank among all the alleles.

### Benchmarking HLA-II ligand predictions

The accuracy of MixMHC2pred was tested in multiple independent HLA-II peptidomics datasets^2,8,30–34^ (Suppl. Table 2). The positives were the peptides of length 12-19 amino acids observed in these samples (removing all the peptides that were also part of the training data from MixMHC2pred for any of the alleles from a given sample). In each sample, we then added 4 times more negatives by randomly sampling human peptides of length 12-19 amino acids.

Predictions with MixMHC2pred and its different variants (see Fig. 2a) were compared with those from NetMHCIIpan (version 3.2^19^ with default parameters) based on the HLA-II typing provided in these studies (Suppl. Table 2). The area under the curve (AUC) of the receiver operating characteristic (ROC) curve was computed for each sample separately (Fig. 2a and Suppl. Fig. 7).

### Benchmarking predictions of epitopes from tetramer assays

All the multimer/tetramer assay data for human CD4+ T cells from the IEDB database^18^ were downloaded (as of 20.07.2018). We then filtered this data to remove peptides with non-standard amino acids and to keep peptides of length 12 and longer that were associated to a known allele based on the “Allele evidence codes”: “MHC binding assay” or “T cell assay -Single MHC type present”. Only interactions involving alleles available in MixMHC2pred were considered.

Predictions with MixMHC2pred and NetMHCIIpan version 3.2^19^ were performed for each peptide with its associated HLA-II allele, both for the positive (1’319 peptides) and the negative (1’040 peptides) cases. The corresponding ROC curve and its AUC are showed in Fig. 2b.

### Selection of candidate viral, bacterial and tumor-associated epitopes

To further benchmark MixMHC2pred, we retrieved a list of known viral, bacterial and melanoma-associated proteins (Suppl. Table 3a) and tested their immunogenicity in two HLA-DRB1*07:01 positive melanoma patients and one HLA-DRB1*07:01 positive healthy donor. Each protein was cut into 20-mers overlapping by 10 amino acids in order to cover all possible 9-mer cores. These 20-mer peptides were then ranked according to the predicted affinity towards HLA-DRB1*07:01 (considering the highest predicted affinity from the 15-mer sub-sequences in each peptide). We then selected the 30 best scoring potential epitopes from the viral/bacterial proteins and the 30 best scoring potential epitopes from the tumor-associated antigens for experimental validation, both for the predictions from MixMHC2pred and NetMHCIIpan.

Matthews correlation coefficients were computed based on the epitopes tested experimentally (e.g. the true negatives from MixMHC2pred are the peptides that had been predicted in the top 60 by NetMHCIIpan but not by MixMHC2pred and that are not immunogenic).

### Peptide synthesis

Peptides were synthesized at the Protein and Peptide Chemistry Facility, University of Lausanne, Switzerland, by standard solid phase chemistry on a multiple peptide synthesiser (Applied Biosystem). All peptides were >90% pure as indicated by analytic HPLC. Lyophilised peptides were diluted in pure DMSO at 10 mg/ml and aliquots at 1 mg/ml in 10% DMSO were prepared and stored at −80°C.

### In vitro peptide stimulation

Peripheral blood mononuclear cells (PBMCs) from two HLA-DRB1*07:01 positive malignant melanoma patients and from one healthy HLA-DRB1*07:01 positive donor were thawed and CD4+ T cells enriched using anti-CD4 microbeads and MiniMACS magnetic separation columns (Miltenyi Biotec, Bergisch Gladbach, Germany). CD4+ T cells were resuspended in RPMI 1640 (Gibco, Dublin, Ireland) supplemented with 2 mM glutamine, 1% (vol/vol) nonessential amino acids, 50 μM 2β-mercaepthanol, penicillin (50 U/ml) and streptomycin (50 μg/ml) (Gibco, Dublin, Ireland), and 8% human serum (Blood transfusion center, Bern) (complete medium) and seeded (0.5×10^6^/well) in 48 well plates to which autologous irradiated (30grey) CD4^−^ T cells were added at a 1:1 ratio. Pools of the selected viral/bacterial or tumor associated peptides (20-mers, Suppl. Table 3b) were added to the wells at a final concentration of 2 μM each. After an overnight period in culture, 500μl of media were replaced by fresh media containing 100IU/ml final of hrIL-2. Every two days the media was refreshed. After 10 days of in vitro expansion, cultures were tested for the presence of antigen-reactive CD4+ T cells. Aliquots of 10^5^ cells were transferred to individual wells of a 96-well plate and stimulated overnight with a mix of multiple peptides distributed in different pools according to a specific matrix (Suppl. Fig. 8b). Brefeldin A at 2.5μg/ml (Sigma-Aldrich, Missouri, USA) was added to each well. A non-stimulated control was added as well as a positive control where cells were stimulated with PMA (Sigma-Aldrich, Missouri, USA) and Ionomycine (Sigma-Aldrich, Missouri, USA) at 50ng/ml and 500ng/ml, respectively. The following day, cells were collected and stained using anti-CD3-APC (clone UCHT1, Beckman Coulter, California, USA) and anti-CD4-FITC antibodies (clone RPA-T4, Biolegend, California, USA), Live/dead feasible Aqua dead cell stain (Invitrogen, California, USA) for 20 minutes at 4°C. Cells were then washed with PBS, fixed and permeabilized using the FOXP3/transcription kit (Invitrogen, California, USA) 30 minutes at room temperature. Finally, the cells were stained for intracellular markers using anti-IFNγ-PE (clone 4SB3, BD, New Jersey, USA) and anti-TNFα-AF700 antibodies (clone Mab11, BD, New Jersey, USA) for 20 minutes at 4°C. Cells were acquired with the CYTOFLEX analyzer (Beckman Coulter) and data were analyzed with Flowjo, LLC software (Oregon, USA). Positive wells for IFNγ and/or TNFα were identified (Suppl. Fig. 8 c-d). Since each individual peptide was only contained in two different pools, by matching the positive wells in the matrix individual immunogenic peptide were selected and evaluated individually. In a similar procedure that was used to evaluate multiple peptides in a matrix format, individual selected peptides were added to newly seeded CD4+ T cells and after an overnight incubation the cells were evaluated for their expression of IFNγ and TNFα using the same method as described above (Suppl. Fig. 8 e-f).

### Patient and neoantigen description

Patient CTE-0007 is a patient with recurrent ovarian cancer. Clinical data and all methodologies for the identification of non-synonymous somatic mutations were already described^35^.

### Identification and validation of neo-epitope specific CD4^+^ TILs

Neoantigen (mutation *D246N* in SGOL1)-specific CD4^+^ TILs were identified in patient CTE-0007 upon co-culture with tandem minigene-transfected autologous B cells. TILs were derived from tumor single cell suspensions and expanded with high-dose IL-2 (Proleukin, 6’000 IU/mL) for 15 days as previously reported^35^. In parallel, CD19^+^ cells were isolated from PBMCs using magnetic beads (Miltenyi) and expanded for 14 days with multimeric-CD40L (Adipogen, Epalinges, Switzerland, 1μg/mL) and IL-4 (Miltenyi, 200 IU/mL). CD40-activated B cells were electroporated using a Neon system (Invitrogen) with 1μg *in vitro* transcribed RNA (Ambion, Foster City, California) coding for 31-mers centered on the specific mutations. Following 16 hours of resting post-electroporation, 10^5^ cells/well of RNA-transfected B cells were co-cultured with TILs at a ratio of 1:1 and incubated over-night in pre-coated ELISpot plates (Mabtech, Nacka Strand, Sweden). Subsequently, T cell activation was validated by intracellular cytokine staining, as described^35^. Either RNA-transfected B cells or B cells loaded with peptides were used as APCs. Neoepitope-reactive CD4^+^ TILs were sorted using a FACSAria IIu, based on CD154 upregulation, as described^36^. CD154-sorted cells were expanded with irradiated feeder cells (PBMCs from two donors) in the presence of OKT3 (Miltenyi, 30 ng/mL) and IL-2 and further interrogated to identify the predicted candidate epitopes by IFN- ELISpot. Additionally, HLA-DR (clone L243, in house production) blocking antibody was added together with cognate peptides. For HLA-restriction analysis, HLA-matched or mismatched CD40-activated B cells were loaded with 2μM of the peptide SPIFKQKKNLRRS for 2 hours before co-culture.

The candidate epitopes have been selected based on the top 5 predictions from MixMHC2pred and NetMHCIIpan among all 13- to 16-mers of the minigene (TDLCFLNSPIFKQKKNLRRSKKRALEVSPAK).

### Code availability

MoDec and MixMHC2pred are freely available as a C++ executable (https://github.com/GfellerLab) for academic non-commercial research purposes.

