## Supplementary Information for "Deep motif deconvolution of HLA-II peptidomes for robust class II epitope predictions"

### Supplementary information for the paper:
### Deep motif deconvolution of HLA-II peptidomes for robust class II epitope predictions

Julien Racle[1,2], Justine Michaux[1,3], Georg Alexander Rockinger[4], Marion Arnaud[1,3], Sara Bobisse[1,3], Chloe Chong[1,3], Philippe Guillaume[1,3], George Coukos[1,3], Alexandre Harari[1,3], Camilla Jandus[4], Michal Bassani-Sternberg[*,1,3] & David Gfeller[*,1,2]

[1]Department of Oncology UNIL CHUV, Ludwig Institute for Cancer Research, University of Lausanne, Epalinges, Switzerland

[2]Swiss Institute of Bioinformatics (SIB), Lausanne, Switzerland

[3]Department of Oncology UNIL CHUV, Ludwig Institute for Cancer Research, University Hospital of Lausanne, Lausanne, Switzerland

[4]Department of Oncology UNIL CHUV, University Hospital of Lausanne, Lausanne, Switzerland

#### Supplementary Notes

##### Further details on MoDec framework

*Definition of the special sum*

The *special sum* "$l \oplus s$" appearing in equation (1) of the log-likelihood equation solved by MoDec is defined in the following way:

$$(l \oplus s)_n = l + s + \left\lceil \frac{\lambda^n - L}{2} \right\rceil - g(s, \lambda^n, L) \qquad \text{(N1)}$$

with

$$g(s, \lambda^n, L) = \begin{cases} 1, & (\lambda^n - L)\ \%\ 2 \neq 0 \ \text{ and } s > 0 \\ \infty, & (\lambda^n - L)\ \%\ 2 \neq 0 \ \text{ and } s = 0 \\ 0, & \text{otherwise} \end{cases} \qquad \text{(N2)}$$

This special sum renders the binding core offsets to be fully symmetric around 0 for each peptide (e.g. only the *s* values -2, -1, 1 and 2 will be defined for a peptide of length 12 when the motif has a length 9, while a peptide of length 13 would additionally have the offset *s=0*).

*Parameter estimation by expectation-maximization*

For a fixed number of motifs $K$, MoDec determines the maximal likelihood from equation (1) through the expectation-maximization algorithm. Similarly to the derivation proposed by Bishop[1], we rewrite equation (1) as

$$\log(\mathcal{L}) = \sum_n \left(W^n \cdot \mathfrak{L}(q^n, \theta, w) + W^n \cdot \mathrm{KL}(q^n \parallel p^n)\right) + \log\left(\mathrm{P}(\theta)\right) \quad \text{(N3)}$$

where $q^n(z^n=(k,s))$ is a normalized distribution defined over some latent variables $z^n$. $q^n$ can also be referred to as the "responsibility of peptide $n$ towards the motif $k$ and binding core offset $s$". The other terms of this equation correspond to:

$$\mathfrak{L}(q^n, \theta, w) = \sum_{z^n} q^n(z^n) \cdot \log\left(\frac{\mathrm{P}(x^n, z^n | \theta, w)}{q^n(z^n)}\right) \quad \text{(N4)}$$

$$\mathrm{KL}(q^n \parallel p^n) = -\sum_{z^n} q^n(z^n) \cdot \log\left(\frac{\mathrm{P}(z^n | x^n, \theta, w)}{q^n(z^n)}\right) \quad \text{(N5)}$$

and

$$p^n = \mathrm{P}(z^n | x^n, \theta, w) \quad \text{(N6)}$$

Note that all $\mathrm{KL}(q^n \parallel p^n)$ correspond to Kullback-Leibler divergences and are always positive or 0.

The expectation-maximization then works briefly as follow:

1. Some initial conditions for the $\theta$ and $w$ are given.
2. "Expectation step": $\theta$, $w$ are fixed and we search for the peptide responsibilities $q^n$ that make the Kullback-Leibler divergences from equation (N5) to be 0.
3. The next step is the "maximization step": the peptide responsibilities are now fixed to those found in previous step and the lower bound from equation (N3) is maximized over $\theta$ and $w$ (i.e. the Kullback-Leibler terms are "removed" as always positives and the other terms are maximized).
4. The steps 2 and 3 are then repeated multiple times until convergence to some maximum.

This algorithm does however not guarantee the convergence towards the global maximum. Therefore, the above iterations starting from different initial conditions are repeated multiple times and the best result is selected.

### Supplementary Figures

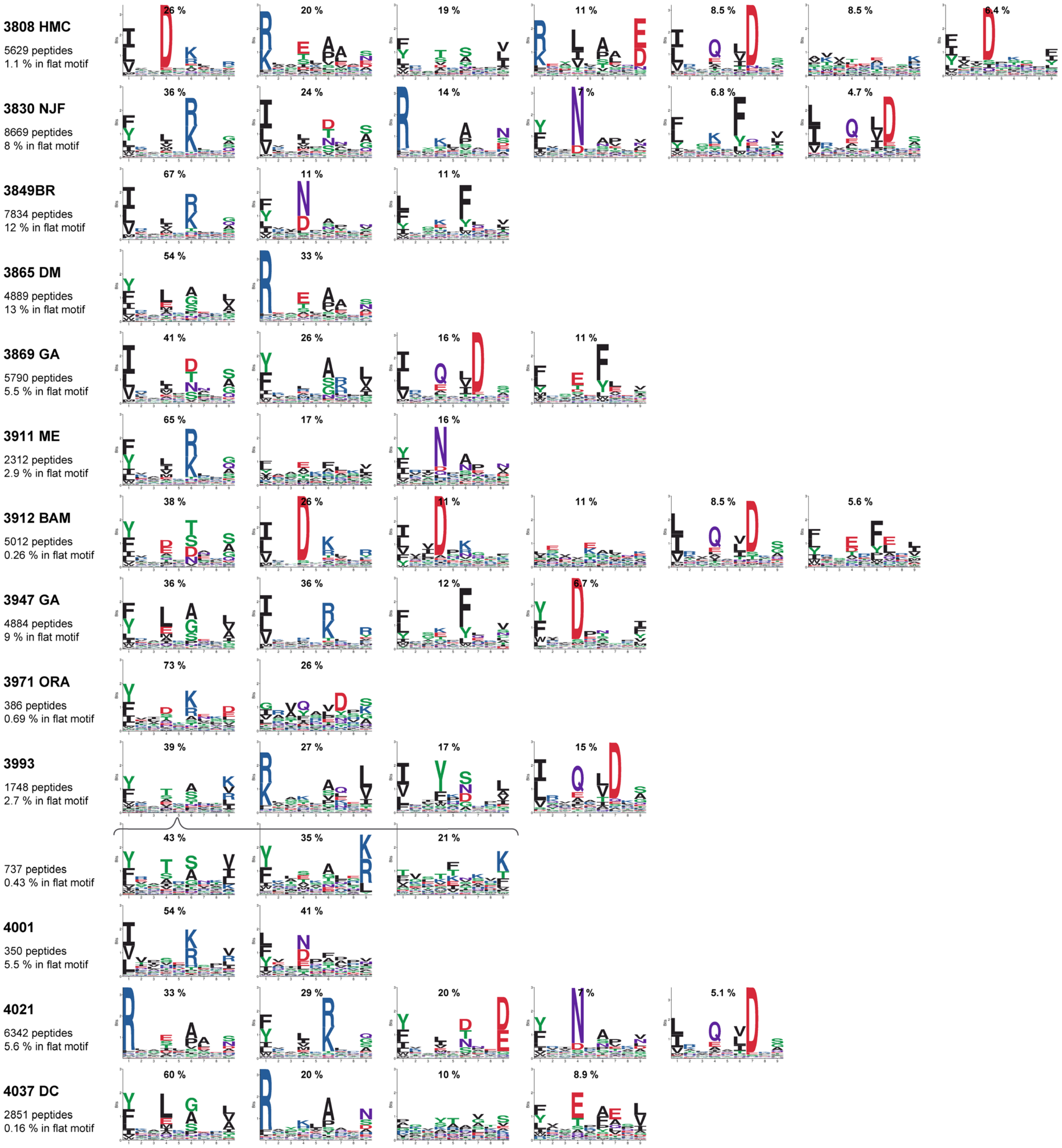


**Supplementary Figure 1 (continued on next page).**

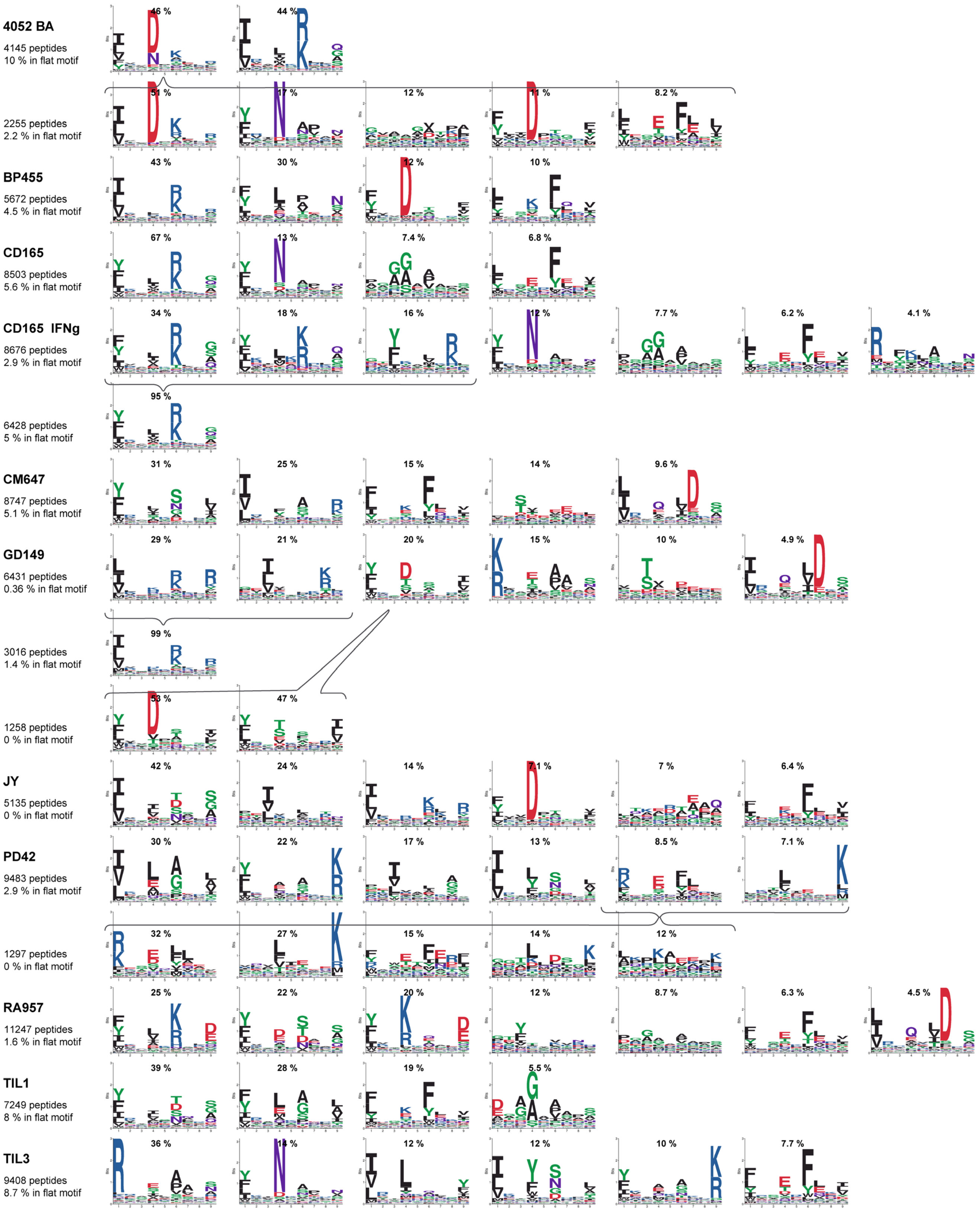


**Supplementary Figure 1. Motifs found by MoDec in the HLA-II peptidomics data.** Each row corresponds to a different deconvolution. Sample name and number of peptides are indicated in the first column as well as the percentage of peptides that best represent false positives (flat motif). Numbers above each motif indicate the percentage of peptides assigned to this motif from the sample. In few cases when a motif seemed to combine multiple specificities, we split it further with MoDec as indicated by the accolades under and above the respective motifs.

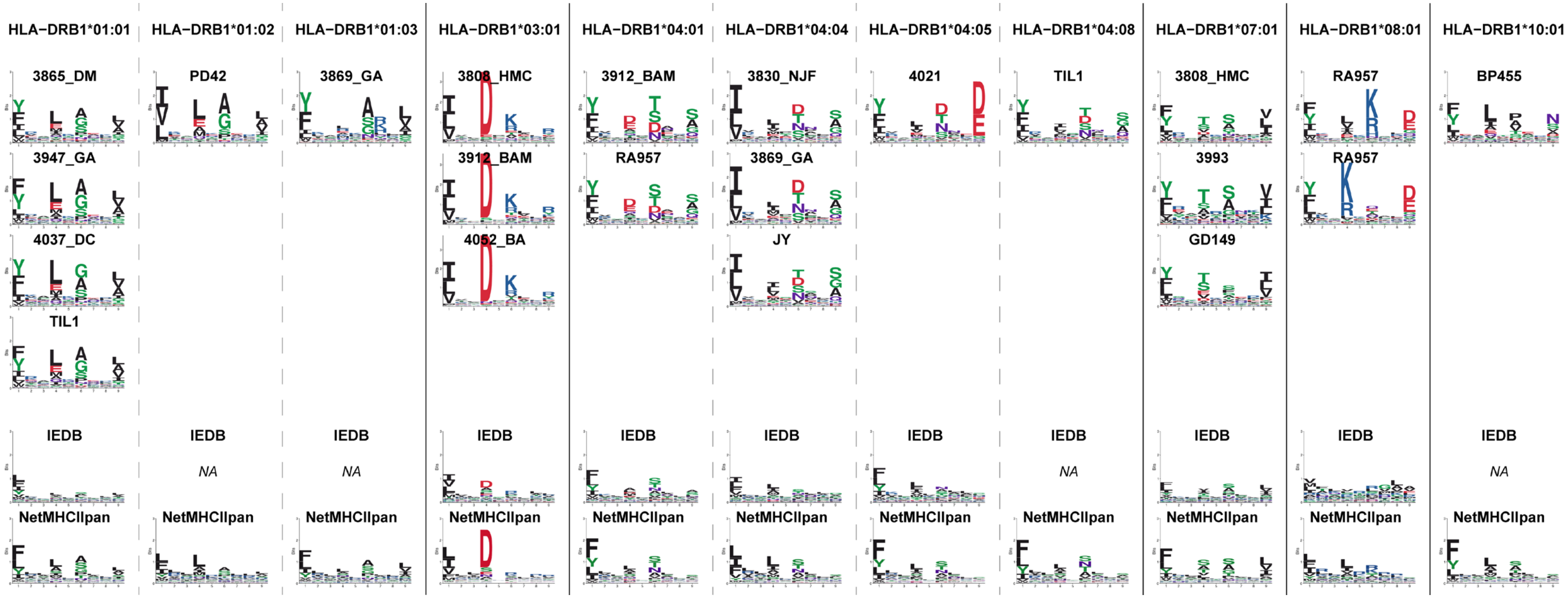


**Supplementary Figure 2 (continued on next page)**

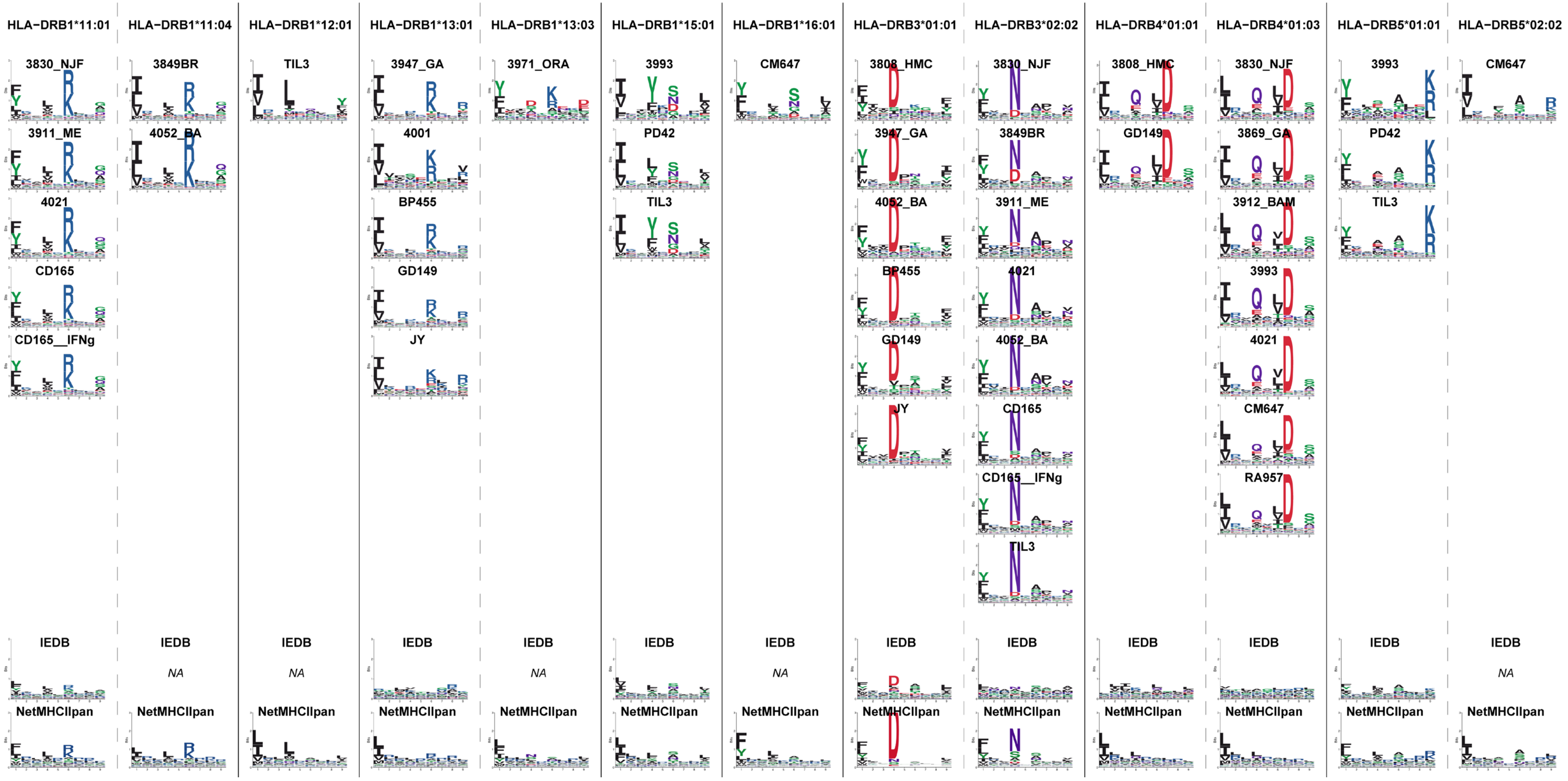


**Supplementary Figure 2 (continued on next page)**

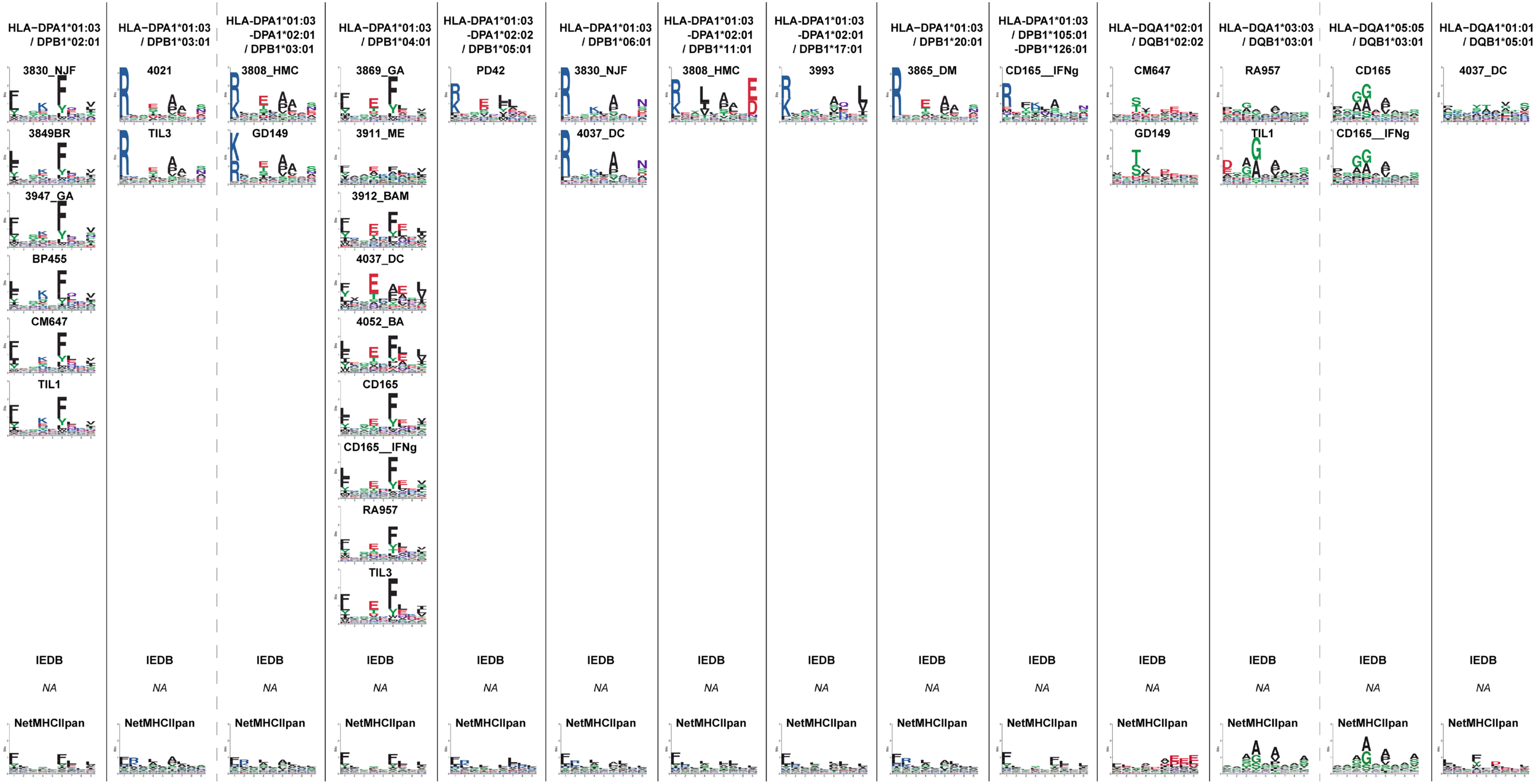


**Supplementary Figure 2. Motifs are highly similar between cell lines and tissues sharing a same allele – the corresponding allele is annotated on top of each column.** The name of the cell line or tissue is indicated above each motif. The last two rows show the motifs obtained from IEDB positive-high data for the respective allele (if such data is available for this allele) and the motif determined from NetMHCIIpan (see Methods). DPA1*02:01 only appears in cell lines and tissues also containing DPA1*03:01, which explains why the two alleles are listed with the corresponding motifs.

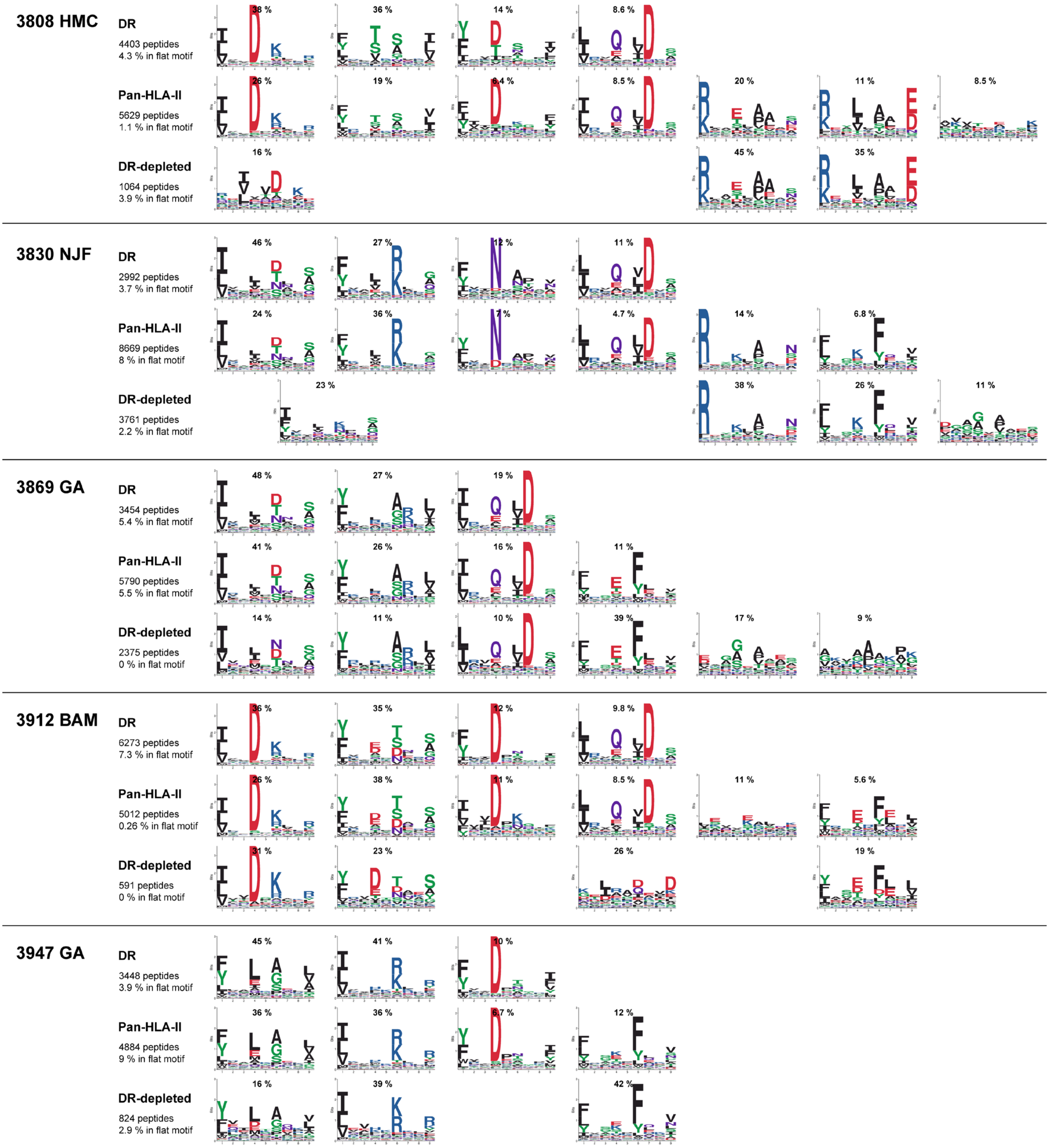


**Supplementary Figure 3 (continued on next page).**

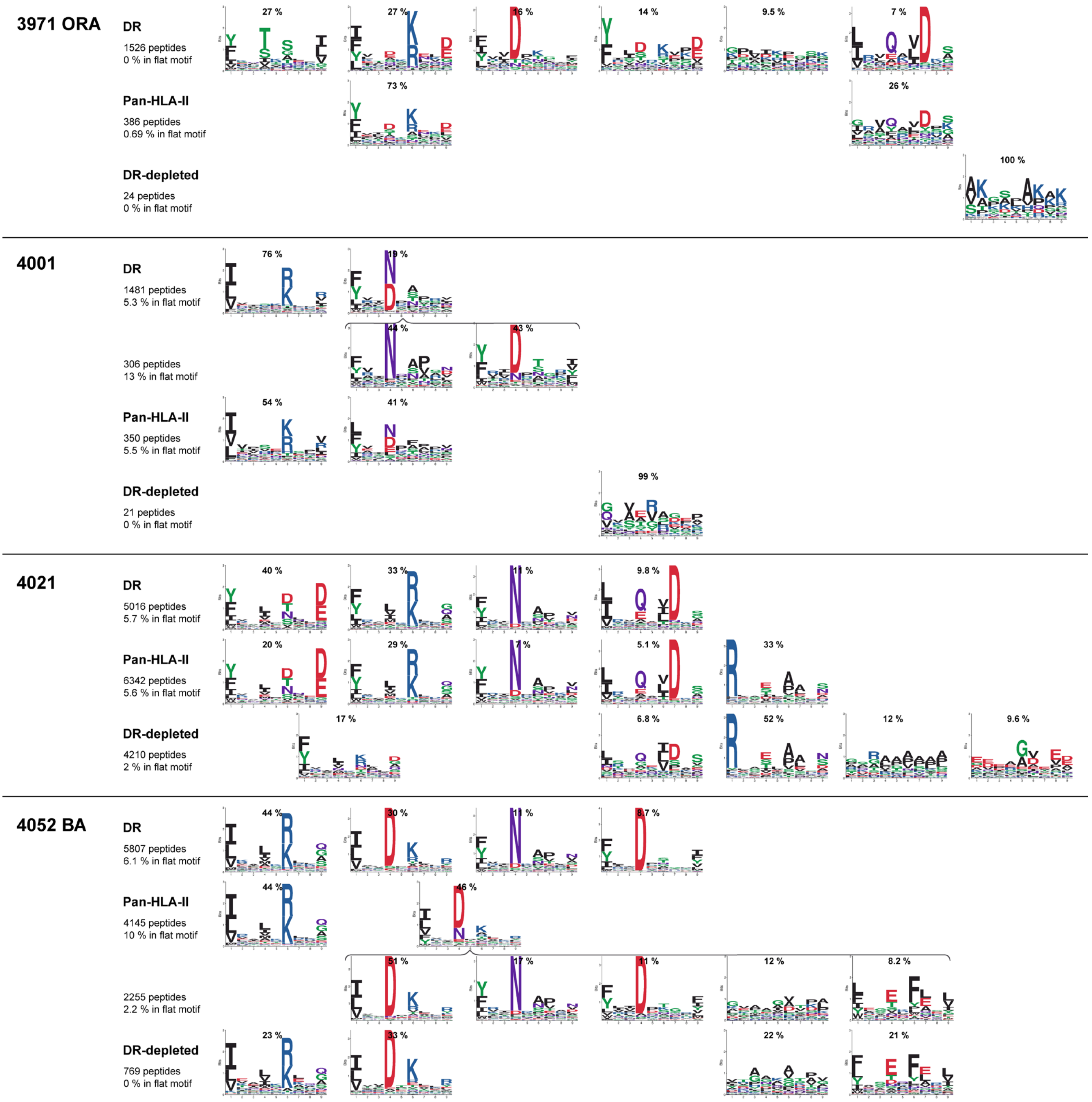


**Supplementary Figure 3 (continued on next page).**

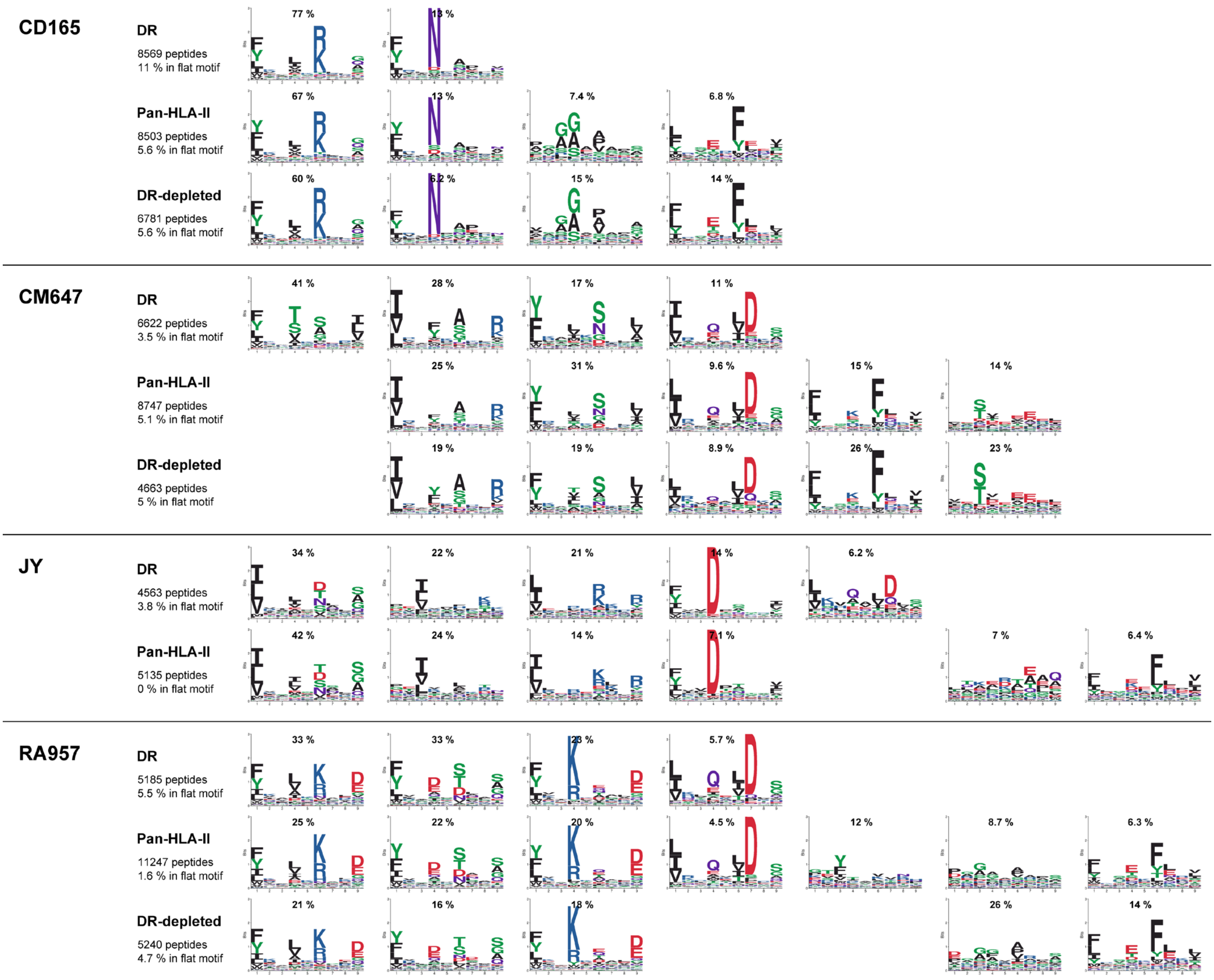


**Supplementary Figure 3. Similar to Supplementary Fig. 1, showing here a comparison of the motifs found with MoDec in HLA-II, HLA-DR and HLA-DR-depleted peptidomics on a same cell line or tissue.**

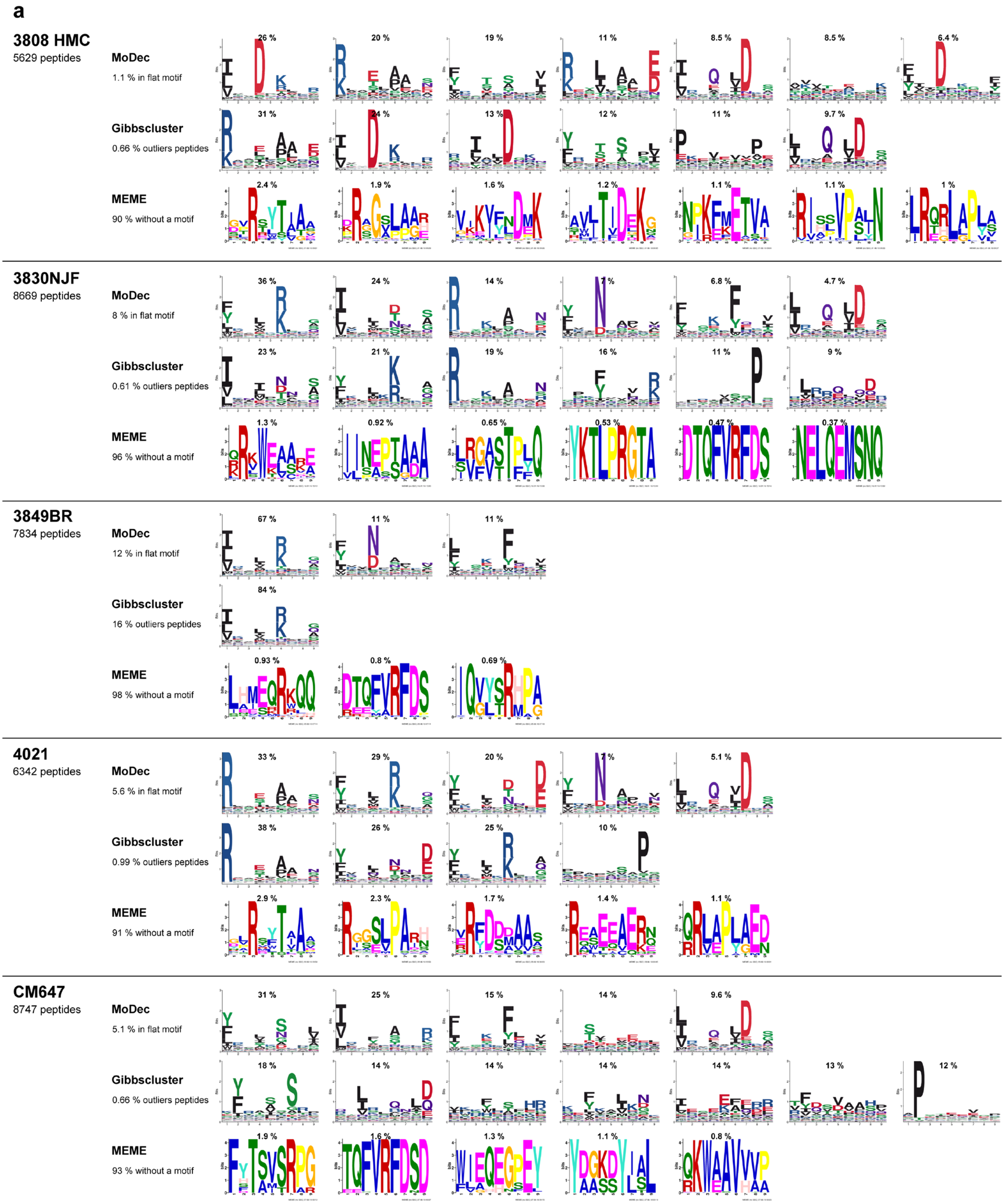


**Supplementary Figure 4 (continued on next page).**

b

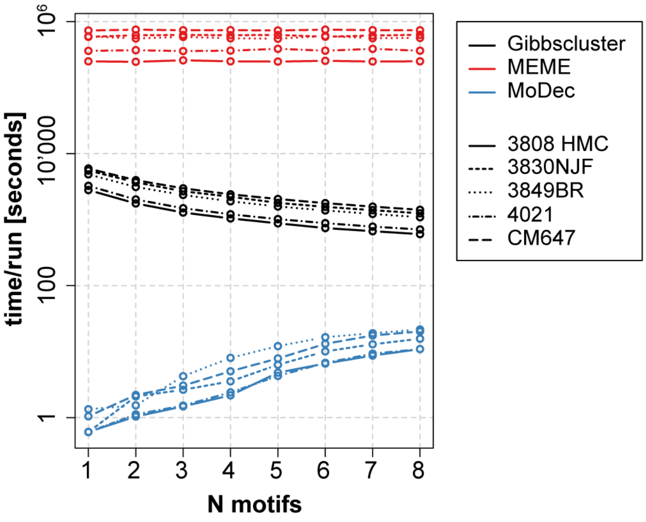


**Supplementary Figure 4. MoDec has a better resolution and is faster than other deconvolution methods. (a)** Comparison of the motifs found by MoDec, Gibbscluster and MEME in 5 different cell lines or tissue samples. MoDec and Gibbscluster both find the motifs containing many peptides, but MoDec finds additional well-defined motifs containing fewer peptides. **(b)** Timing comparison of the 3 methods in function of the number of motifs searched.

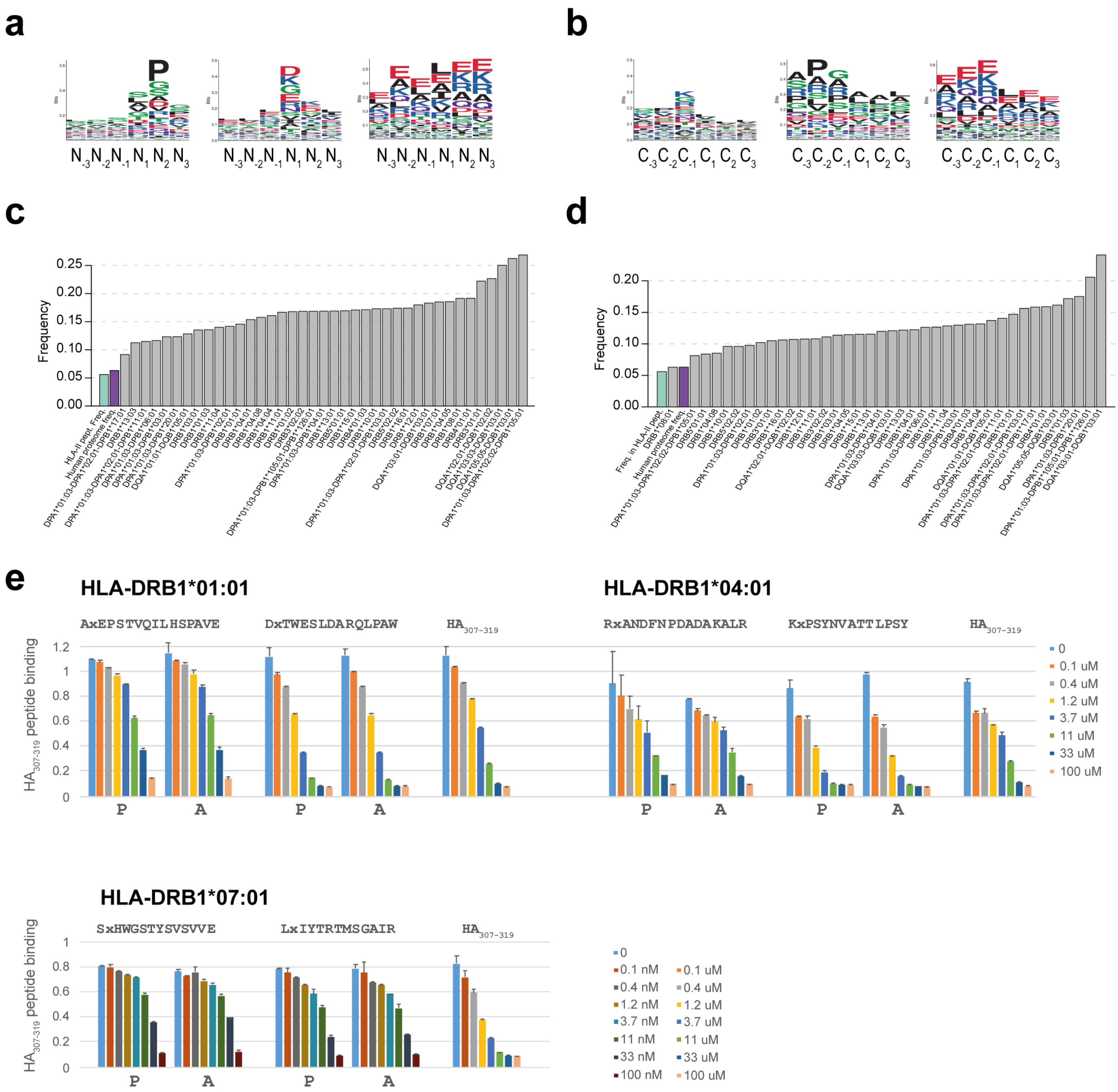


**Supplementary Figure 5. Characteristics of the N- / C-terminus region of the peptides. (a-b)** Motifs determined by MoDec for the three amino acids upstream and downstream of the N-terminus (a) or C-terminus (b) when grouping all HLA-II peptidomics data. **(c-d)** Frequency of proline at the 2nd position downstream of N-terminus (c) or 2nd position upstream of C-terminus (d) of the peptide in function of the allele to which the peptide is associated. Colored bars represent the proline frequency in the human proteome and the proline frequency at any position in HLA-II peptidomics. **(e)** Competition assays for different alleles between $HA_{307\text{-}319}$ and various peptide variants containing proline at the 2nd position or alanine (the “x” in the peptide sequences is replaced by proline or alanine as indicated below the bars; $HA_{307-319}$ is used as a control).

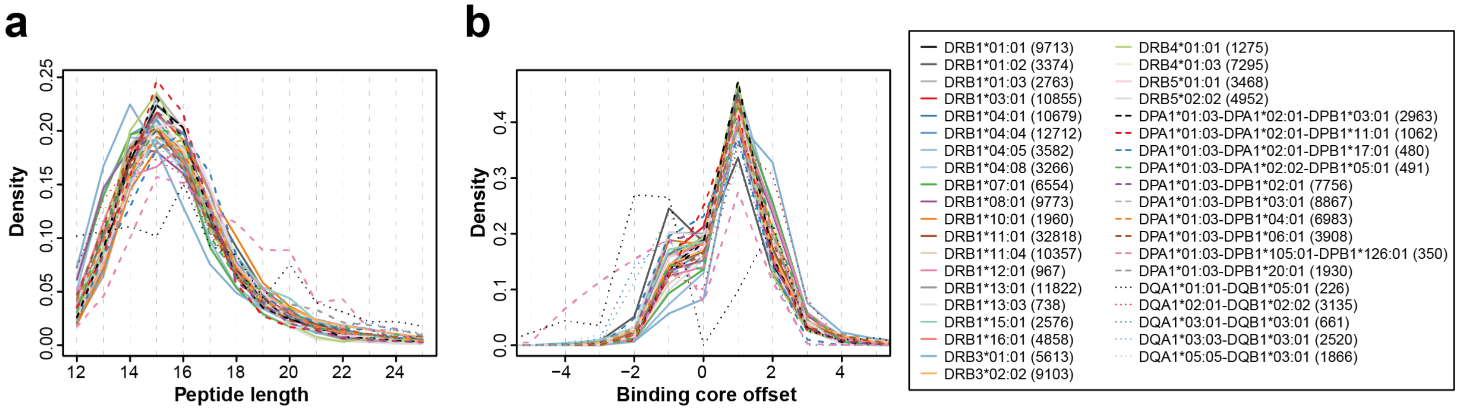


**Supplementary Figure 6. Peptide length (a) and binding core offset (b) are independent of the allele.** Number of peptides associated to each allele is indicated in parenthesis. The outliers' distributions are those from alleles with too few peptides for robust statistics.

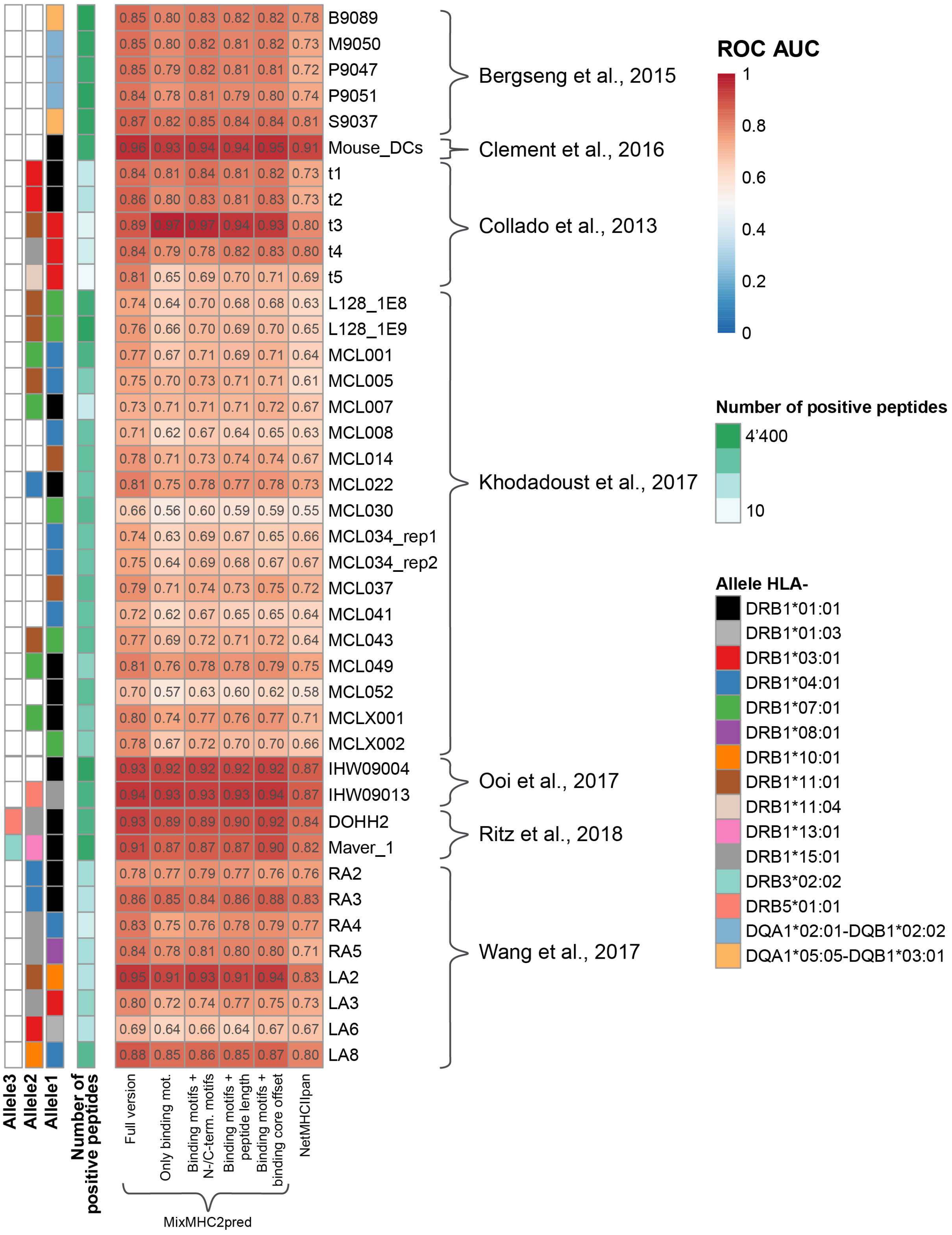


**Supplementary Figure 7. Detailed results from the comparison of the predictors on independent HLA-II ligand data, showing the ROC AUC from each sample.**

a

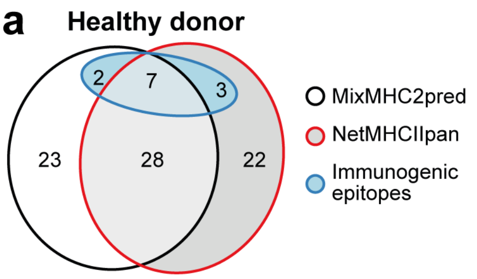


b

| Matrix | A1 | A2 | A3 | A4 | A5 | A6 | A7 |
|---|---|---|---|---|---|---|---|
| B1 | P1 | P2 | P3 | P4 | P5 | P6 | P7 |
| B2 | P8 | P9 | P10 | P11 | P12 | P13 | P14 |
| B3 | P15 | P16 | P17 | P18 | P19 | P20 | P21 |
| B4 | P22 | P23 | P24 | P25 | P26 | P27 | P28 |
| B5 | P29 | P30 | P31 | P32 | P33 | P34 | P35 |
| B6 | P36 | P37 | P38 | P39 | P40 | P41 | |

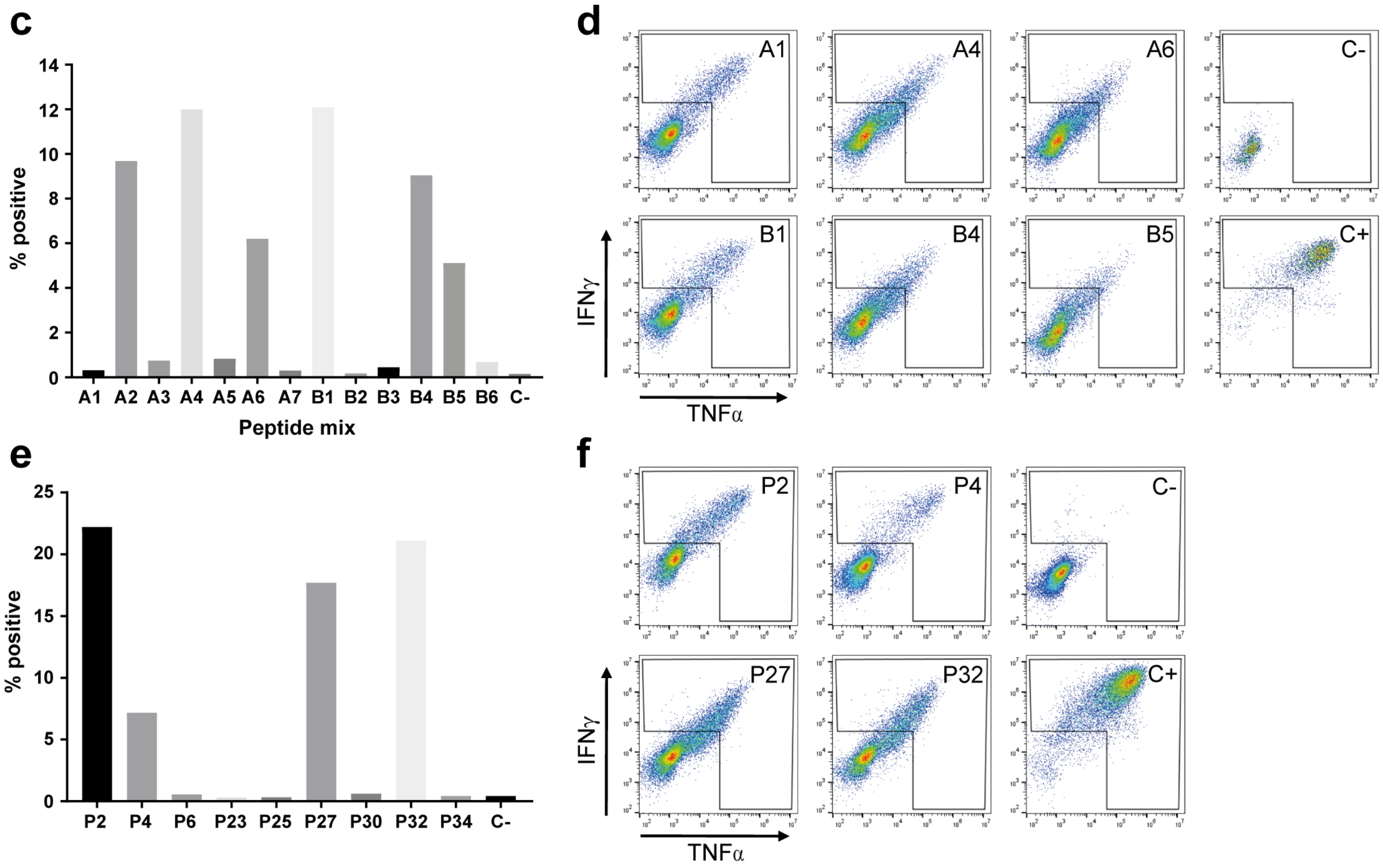


**Supplementary Figure 8. Determining class II immunogenic epitopes from viral, bacterial and tumor-associated antigens. (a)** Similar to Fig. 2c with the results from a healthy donor (MCC of 0.039 for MixMHC2pred and 0.11 for NetMHCIIpan). **(b-f)** Representative example of the immunogenicity screening procedure, showing the results from the viral/bacterial epitopes of the melanoma patient LAU 1352. (b) Matrix used for the screening of all viral/bacterial peptides. A-labelled mixes from 1 to 7 contain vertically listed peptides and B-labelled mixes from 1 to 6 contain horizontally listed peptides. (c) Readout in % positive cells (positive for intracellular IFNγ and TNFα) following stimulation with the peptide matrix mixes. (d) Dot plot representation of the wells positive for both IFNγ and TNFα, and of the negative (C-) and positive controls (C+). (e) Readout of the single peptides tested individually, peptides that were selected based on the positive wells from the peptide mixes stimulation. (f) Dot plot representation of the wells reacting to individual peptides and the positive and negative controls.

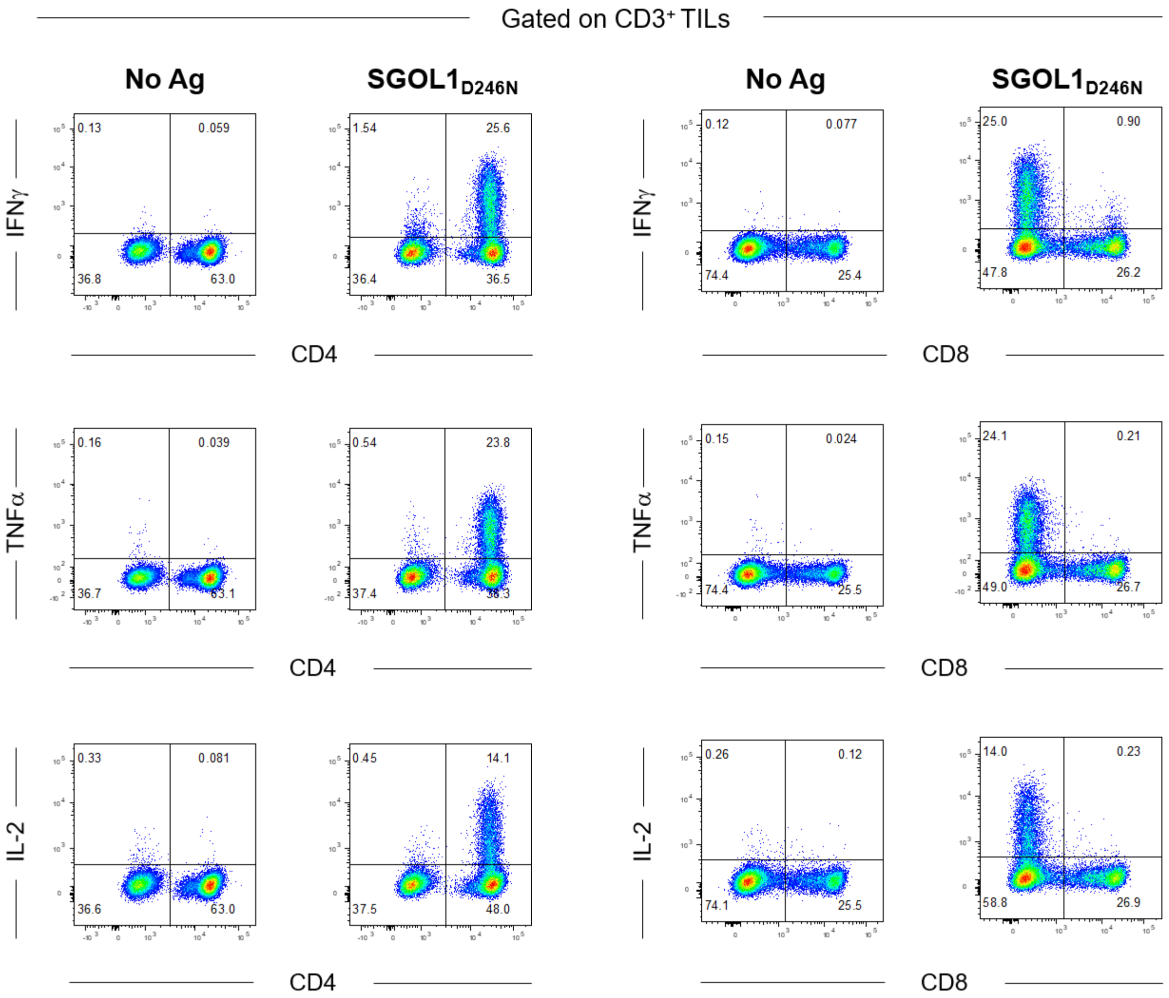


**Supplementary Figure 9.** ***SGOL1**$_{D246N}$***-specific tumor infiltrating lymphocytes from patient CTE-0007 are CD4+ T cells.** Intracellular cytokine staining showing the frequency of viable IFNγ, TNFα, and IL-2 cytokine- producing CD4+ TILs following stimulation with the 31-mer peptide containing the identified neoepitope from *SGOL1*$_{D246N}$ gene.

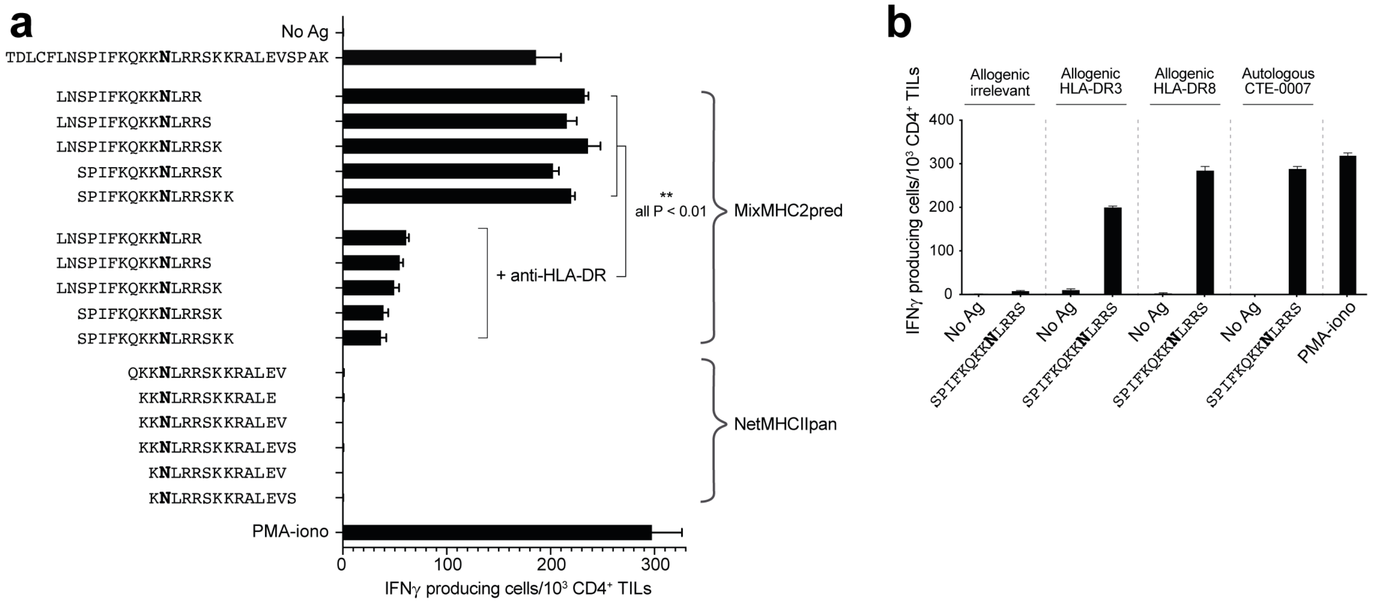


**Supplementary Figure 10. Validation of neoepitope and HLA restriction. (a)** Neoepitope-specific CD4+ TILs from patient CTE-0007 were isolated based on CD154 upregulation following stimulation with the 31-mer peptide encompassing the neoepitope ($SGOL1_{D246N}$). Neoepitope-specific CD4+ TILs were interrogated by IFNγ ELISpot with the candidate epitopes from MixMHC2pred and NetMHCIIpan (mutation indicated in bold in the peptide sequences). Autologous CD40-activated B cells were used as APCs. T cell stimulation was inhibited in the presence of anti-HLA-DR antibody (clone L243). **(b)** Neoepitope specific CD4+ TILs from patient CTE-0007 were re-challenged with CD40-activated B cells pulsed with 2μM of the peptide SPIFKQKK**N**LRRS. Autologous B cells, allogenic B cells sharing HLA-DRB1-03:01 (DR3) or HLA-DRB1-08:01 (DR8) alleles and irrelevant allogenic B cells (neither DR3 nor DR8) were used as APCs. Screening by IFNγ ELISpot (*PMA-iono*: phorbol-myristate-acetate + ionomycin, *No Ag*: no peptide).

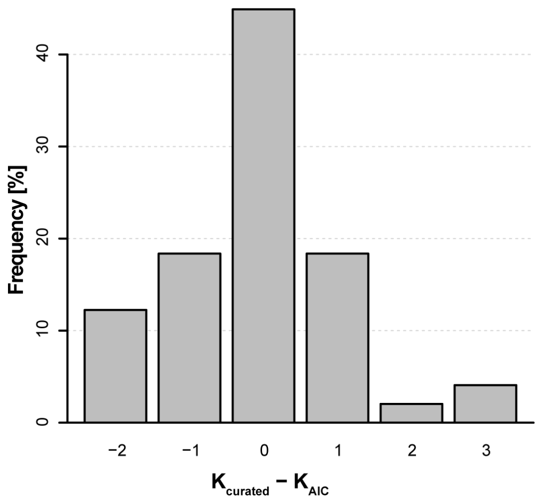


**Supplementary Figure 11. Comparison between the number of motifs determined by the AIC and after manual curation.** The best number of motifs is most often directly selected by the AIC.

#### Supplementary Tables and Supplementary Data

The Supplementary Tables and Supplementary Data will be available in the peer reviewed version of this manuscript.
